# The axon guidance cue SEMA3A promotes the aggressive phenotype of basal-like PDAC

**DOI:** 10.1101/2023.02.25.529923

**Authors:** Francesca Lupo, Francesco Pezzini, Elena Fiorini, Annalisa Adamo, Lisa Veghini, Michele Bevere, Cristina Frusteri, Pietro Delfino, Sabrina D’Agosto, Silvia Andreani, Geny Piro, Antonia Malinova, Francesco De Sanctis, Davide Pasini, Rita T. Lawlor, Chang-il Hwang, Carmine Carbone, Ivano Amelio, Peter Bailey, Vincenzo Bronte, David Tuveson, Aldo Scarpa, Stefano Ugel, Vincenzo Corbo

## Abstract

Pancreatic ductal adenocarcinoma (PDAC) is a lethal disease with few available therapeutic options. Two transcriptional cancer cell states have been consistently reported in PDAC, with the basal-like/squamous phenotype displaying a more aggressive biological behavior. Genetic and epigenetic dysregulation of the axon guidance pathway are common in PDAC, yet our understanding of its biological relevance is limited. Here, we investigated the functional role of the axon guidance cue SEMA3A in sustaining the progression of PDAC. We integrated available transcriptomic datasets of human PDAC with *in situ* hybridization analyses of patients’ tissues to find that SEMA3A is expressed by stromal cells and selectively enriched in epithelial cells of the basal-like/squamous subtype. We found that both cell-intrinsic and cell extrinsic factors instructing the basal-like/squamous subtype induce expression of SEMA3A in PDAC cells. *In vitro*, SEMA3A promoted cell migration as well as anoikis resistance. At molecular level, these phenotypes were associated with increased FAK signaling and enrichment of gene programs related to cytoskeleton remodeling. Accordingly, SEMA3A provided mouse PDAC cells with greater metastatic competence. In mouse orthotopic allografts, SEMA3A remodeled the TME by favoring infiltration of tumor-associated macrophages and exclusion of T cells. Mechanistically, SEMA3A functioned as chemoattractant for macrophages and favored their polarization towards an M2-like phenotype. In SEMA3A^high^ tumors, depletion of macrophages resulted in greater intratumor infiltration by CD8+ T cells and increased sensitivity of these tumors to chemotherapy. Overall, we show that SEMA3A contributes to the malignant phenotype of basal-like PDAC.

## Introduction

Pancreatic ductal adenocarcinoma (PDAC) is a malignancy of the exocrine pancreas and the deadliest cancer worldwide [1]. Most patients present with an unresectable disease at diagnosis that is treated with chemotherapy-based regimens [2]. Overall, PDAC is poorly responsive to systemic treatments [2], including immune-oncology approaches. Evidence from studies addressing recurrences of PDAC following radical surgery suggests that pancreatic cancer is a systemic disease at presentation and its biology, rather than the surgical procedures, determines patterns of recurrence [3, 4, 5]. As it stands, understanding the mechanisms of tumor progression and dissemination in PDAC is vital to improve patients’ outcomes in the long-term. At histopathological level, PDAC tissues are characterized by a prominent stromal reaction with abundant cancer-associated fibroblasts (CAFs) and macrophages, while T cells are usually excluded. Macrophages represent the prevalent leukocyte population in the PDAC stroma and contribute to shaping a highly immunosuppressive microenvironment [6]. Expression profiles analyses have evidenced two main subtypes of PDAC cells, which show distinct biological properties [7, 8, 9, 10]. The basal-like/squamous subtype is characterized by the loss of pancreatic endodermal identity and shows the more aggressive biological behavior [7, 10]. Accordingly, basal-like/squamous cells accumulate in advanced stages of the disease [8]. Preclinical studies showed that PDAC molecular subtypes are not permanently encoded, i.e., “molecular subtype switching” is observed in response to microenvironmental changes [6, 11]. Indeed, these cancer cell states results from the integration of cell autonomous and non-cell autonomous inputs. The basal-like/ squamous subtype is characterized by the enrichment for inactivation of *TP53* and chromatin modifiers as well as by p63/ΔNp63 driven genetic programs [7, 8, 10]. Nonetheless, different type of microenvironmental pressures can lead to the emergence of the basal-like phenotype [11] and accordingly molecular signatures indicative of a challenging microenvironment (e.g, hypoxia, fibrosis) represent core gene programmes of this subtype [7, 8]. Genetic and epigenetic dysregulation of the Axon guidance pathway have been repeatedly reported in PDAC [12, 13, 14]. Recently, Krebs and colleagues showed the enrichment of axon guidance-associated gene sets in basal-like as well as high grade PDAC [15]. Furthermore, neuronal-like progenitor cell states have been reported in undifferentiated tumors [16] and positively selected in post-treatment tumors [17]. Most of the previous studies have focused on investigating the role of members of the Slit/Robo axis on the PDAC malignant traits of PDAC as well as its cell identity [15, 18, 19, 20, 21]. Semaphorins are the largest family of axon guidance cues, which were originally identified as chemorepellent proteins in the nervous system [22, 23]. Semaphorins can be secreted, GPI-anchored or transmembrane proteins, and have now recognized functions outside the central nervous system, including angiogenesis and regulation of immune responses [24, 25]. SEMA3A is a class 3 Semaphorins, i.e., secreted, whose elevated tissue expression is a negative prognostic marker in PDAC [12, 13]. Nonetheless, the functional role of Semaphorins in PDAC remains to be elucidated. Here, we investigated whether the Semaphorins signaling pathway contributes to shaping aggressive PDAC phenotypes. Integrating bulk and single-cell RNA-sequencing data with *in situ analysis* of PDAC tissues, we demonstrated that *SEMA3A* expression is prominent in the stroma of PDAC and further enriched in the epithelial cells of the basal-like/squamous subtype. We found that both cell-intrinsic and cell extrinsic factors promoting the basal-like/squamous subtype induce expression of *SEMA3A* in PDAC cells. Mechanistically, SEMA3A acts cell autonomously to promote mesenchymal-like traits, including anoikis resistance, through the activation of Focal adhesion kinase (FAK). *In vivo*, SEMA3A promotes the intratumor infiltration of macrophages that mediates T cell exclusion. Finally, depletion of macrophages with a CSF1R monoclonal antibody restored the sensitivity of SEMA3A expressing tumors to gemcitabine.

## RESULTS

### The expression of class 3 Semaphorins is associated with the basal-like/ squamous phenotype of PDAC

We interrogated three distinct PDAC transcriptomic datasets [7, 8, 26] to investigate whether expression of individual Semaphorins was differently linked to the two reported PDAC subtypes. The expression level of 4 Semaphorins significantly discriminated basal-like from classical tumors in the ICGC[7] and the PanCuRx[8] cohorts (**Figure 1A** and **Figure S1A**). In particular, the expression of *SEMA3A, SEMA3C*, and *SEMA3F* was significantly enriched in the basal-like tumors of both the ICGC and the PanCuRx cohorts, while *SEMA4B* levels were enriched in the classical tumors of the same datasets. Basal-like/squamous PDAC is instructed by both cell-intrinsic and cell-extrinsic factors [7, 8, 10, 11]. First, we explored the correlation between Semaphorins’ expression and gene programmes previously linked to this subtype in 3 transcriptional datasets[7, 8, 26] (**Figure 1B and Figure S1B**). Of the interrogated Semaphorins, *SEMA3A* and *SEMA3C* showed the highest correlation with basal-like/squamous transcriptional signatures, including those indicative of a challenging microenvironment (e.g., hypoxia and fibrosis) (Figure 1B and Figure S1B). As expected, levels of *SEMA4B* were negatively correlated with basal-like/squamous gene sets (Figure 1B and Figure S1B). Therefore, we decided to focus on SEMA3A and SEMA3C. In keeping with the correlative analyses, we identified higher SEMA3A expression in whole cell lysates from cells with prominent squamous features including expression of TP63/ΔNp63 (Colo357, L3.6pl, BxPC3) or CK5 (Hs766t, see D’Agosto et al. [27]) (**Figure 1C**). SEMA3C showed a more promiscuous pattern of expression in human cell lines (**Figure S1C**). Transient downregulation of *p63* was sufficient to reduce SEMA3A but not SEMA3C expression in Colo357 and L3.6pl cell lines (**Figure 1D and Figure S1D**). Therefore, in established human PDAC cell lines the expression of SEMA3A correlates with basal-like/squamous features and can be modulated by perturbation of p63. To assess whether dysregulation of *Sema3a* and *Sema3c* is associated with early neoplastic transformation, we evaluated their expression in mouse organoids established from normal (mN), preneoplastic (PanIN, mP), neoplastic (mT), and metastatic (mM) tissues. Overall, the expression levels of both *Sema3a* and *Sema3c* was variable among stage-matched cultures. However, we found a trend towards an increase in *Sema3a* expression in advanced stage cultures, with a significant difference between mM and mN cultures (**Figure S1E**). Accordingly, we observed a significant increase in SEMA3A protein expression and secretion in mM, but not in mT, as opposed to cultures of normal pancreatic cells (**Figure S1E**). mT organoids established from KPC mice, differently from mM organoids, retain the wild-type copy of *Trp53* [28, 29]. The *in vivo* progression towards invasive tumors is almost invariably associated with loss of heterozygosity (LOH) of *Trp53* [28, 29, 30, 31], which has been recently to show to lead to recurrent copy-number alterations including gains of chromosomal regions containing several Semaphorins on chromosome 5 [31]. Therefore, we explored whether expression of SEMA3A/3C was affected by the p53 status *in vitro*. We treated early-passage mT with 10 µM of Nutlin-3A, which has been reported to deplete *Trp53* wild-type cells [32] (**Figure S1G**). Loss of the wild-type copy of *Trp53* in mT was associated with increased transcriptional and protein expression of SEMA3A while levels of SEMA3C were unaffected (**Figure 1E** and **Figure S1H**). Furthermore, only in cells displaying LOH of *Trp53* we could observe a significant induction of *Sema3a* expression following forced expression of p63. Overall, these results suggest that the p53 mutation combined with loss of heterozygosity promote the expression of *Sema3a* but not *Sema3c* in mouse PDAC cells. Furthermore, in line with the human data, forced expression of p63 in KPC monolayer cell cultures led to increased expression of *Sema3a*, which was associated with increased occupancy of its promoter by the transcription factor (**Figure 1F, Figure S1I**). Conversely, transient downregulation of mutant *Kras* in KPC cell lines did not lead to changes in *Sema3a* expression while reducing the levels of genes downstream of mutant Kras signaling such as *Nq01* and *Sema3c* [33, 34] (**Figure S1J**).

**Figure 1.**
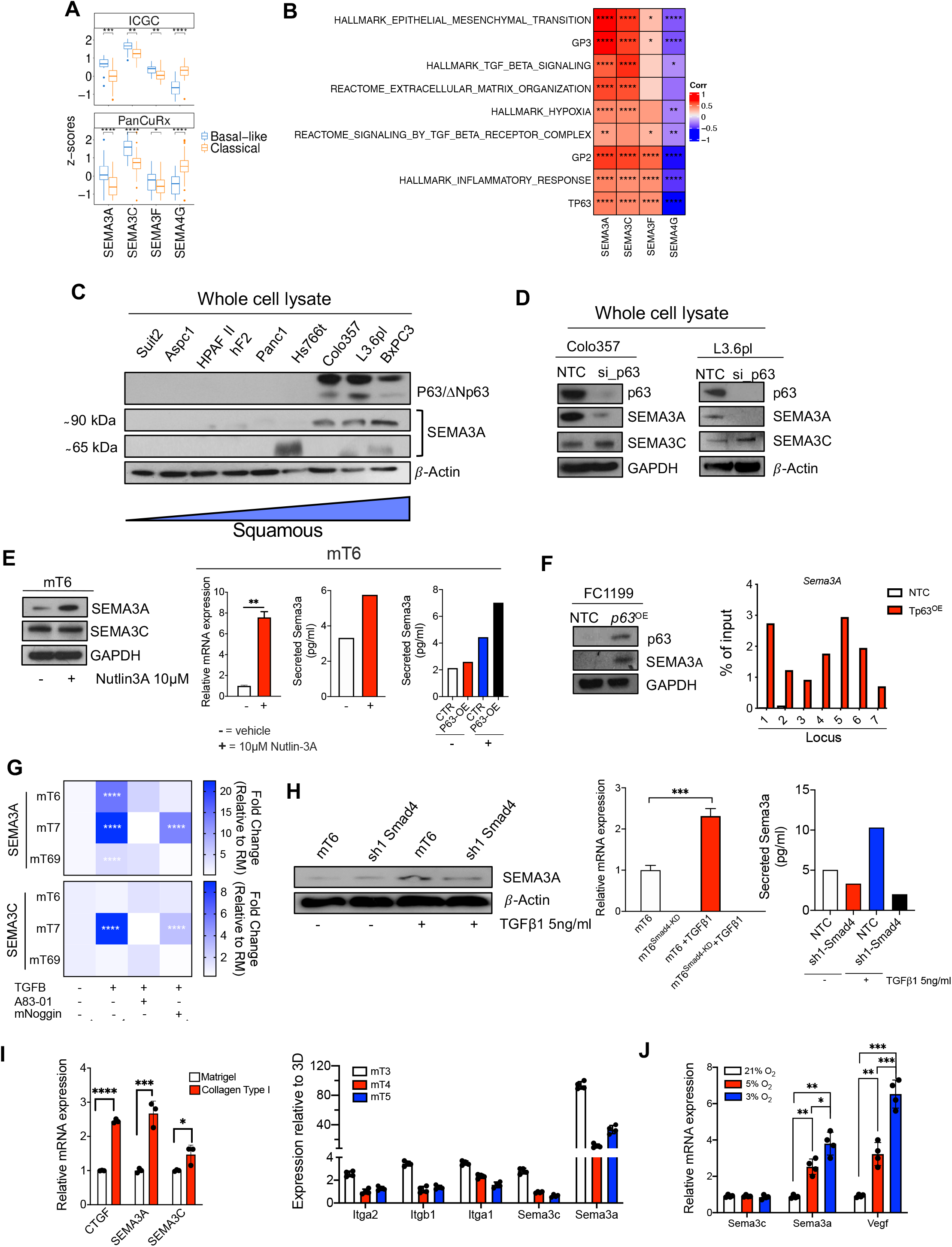
Cell intrinsic and cell extrinsic inputs eliciting SEMA3A expression in PDAC cells. **A**. Boxplot of *SEMA3A, SEMA3C, SEMA3F*, and *SEMA4B* Z-scores stratified by the Moffitt subtypes [10] in the ICGC[7], and the PanCuRx[8] cohorts. **, p < 0.01; ***, p < 0.001; and ****, p < 0.0001 by Student t test. **B**. Heatmap showing correlation (Spearman’s correlation) between the indicated Semaphorins and basal-like/ squamous associated gene programs in the ICGC cohort. GP2 and GP3 refers to the core gene programmes defining the squamous subtype in Bailey et al. [7] All annotated boxes, p < 0.05. **C**. Immunoblot analysis of p63 and SEMA3A in whole cell lysates of different human pancreatic cancer cell lines order based on their basal-like/squamous identity[7, 10] as assessed using GSVA, ssgsea method, on RNA-Seq data. Beta actin was used as loading control. **D**. Immunoblot analysis of p63, SEMA3A, and SEMA3C in whole cell lysates from Colo357 (left) and L3.6pl (right) squamous cell lines transfected with either non-targeting control (NTC) of siRNA targeting p63. β-actin or GAPDH were used as loading control. **E**. Left panel, immunoblot analysis of SEMA3A and SEMA3C in whole cell lysates from mT6 treated with vehicle or Nutlin-3A (see methods). GAPDH was used as loading control. Changes in the expression (qPCR) or secretion (ELISA) of SEMA3A were detected in mT6 following Nutlin-3A treatment (right panel). **F**. Immunoblot analysis of p63 and SEMA3A in KPC 2D cell lines (FC1199) transduced with either an empty vector (NTC) or a p63 ORF. On the right, anti-p63 ChIP-qPCR analysis of seven different genomic regions upstream of the promoter of *Sema3a*. The ChIP-qPCR signal of each sample was normalized to its own input. **G**. Heatmap showing changes in the expression of *Sema3a* (top) and *Sema3c* (bottom) relative to the reduced media condition (RM, without A83-01 and mNoggin) in 3 different tumor organoid cultures treated as indicated. Data are mean of three technical replicates. ****, p < 0.0001 by unpaired Student t test. **H**. Immunoblot analysis of SEMA3A in whole cell lysates from SMAD4 proficient and deficient mT6 organoids that were treated with either vehicle or TGF-β1 for 48 hours. β-actin was used as loading control (left panel). qPCR analysis (middle panel) and ELISA of SEMA3a in mT6 organoids treated as shown in the left panel. ***, p < 0.001 by Student t test. **I**. On the left, changes in the expression of Ctgf, Sema3a, and Sema3c in mouse tumor organoids grown embedded in collagen type I for 48 hours. On the right, changes in the expression levels of Itga2, Itgb1, Itga1, Sema3c, and Sema3a in mouse tumor organoids adapted to grow on plastic. Data are represented as mean value ± S.D (n= 3 technical replicates). *, p < 0.05; ***, p < 0.001; and ****, p < 0.0001 by Student t test. **J**. Changes in the expression levels of *Sema3c, Sema3a*, and *Vegf* in mouse tumor organoids cultivated under different O_2_ concentration for 24 hours. Results are shown as mean ± S.D. of 4 independent experiments. ***, p < 0.001; **, p < 0.01; and *, p < 0.05 by Student t test.

### Environmental cues induce the expression of SEMA3A in mouse PDAC cells

Next, we investigated the effect of microenvironmental factors on Sema3a/3c expression. Elevated TGF-β signaling as well as ECM-related gene programs are enriched in basal-like tumors [7]. The organoid culture medium contains TGF-β inhibitors (A83-01 and mNoggin) yet their removal from the media did not alter the expression of both Sema3a and Sema3c (data not shown). To test whether stromal-derived TGF-β could elicit Sema3a/3c expression in organoids, we treated three different mT cultures with recombinant TGF-β1. The treatment invariably led to increased *Sema3a* expression (**Figure 1G**) while we observed a context-dependent effects for Sema3c (Figure 1G). The stimulatory effect of TGF-β1 on *Sema3a* expression could be blocked by the downregulation of SMAD4 (**Figure 1H** and **Figure S1K**). Then, we tested the effect of matrix rigidity on Sema3a/3c expression by growing mouse tumor organoids on substrates displaying different stiffness for 48 hours. The cultivation of mT in 3 mg/mL of collagen type I significantly induced the expression of the mechanosensitive gene *Ctgf* as well as of Sema3a and Sema3c although to a different extent (**Figure 1I**). Furthermore, the expression of *Sema3a*, but not of *Sema3c* and several Integrins, was significantly induced when mT organoids were adapted to grow on plastic (Figure 1I). Hypoxia-related gene programs are also significantly enriched in basal-like/squamous PDAC [7]. Lowering the concentration of O_2_ significantly induced a dose-dependent expression of the hypoxia-responsive gene *Vegf*, of *Sema3a*, but not of *Sema3c* in KPC cell lines (**Figure 1J**). Altogether, our results show that *Sema3a* is responsive to both cell intrinsic and cell extrinsic inputs that define aggressive PDAC phenotypes. These findings prompted us to investigate whether and how SEMA3A contributes to shape aggressive PDAC phenotypes.

### SEMA3A expression in normal and malignant pancreatic tissues

Elevated tissue expression of SEMA3A has been previously linked with dismal outcomes in PDAC [12]. Herein, we sought to identify the cell types expressing SEMA3A within pancreatic tissues. First, we interrogated publicly available scRNA-Seq data from normal pancreatic tissues [35, 36, 37, 38] to explore *SEMA3A* expression in the different tissue compartments. Only few epithelial cells (150/6321) had detectable levels of *SEMA3A* and were mostly neuroendocrine cells (**Figure 2A**). In situ hybridization (ISH) analysis of normal pancreatic tissues confirmed the scRNA-Seq data (**Figure 2B**) and further showed very rare cells expressing *PLXNA1*, the receptor for SEMA3A. Next, we explored scRNA-Seq data of human PDAC tissues [8, 39, 40, 41] to find that expression of *SEMA3A* is not restricted to epithelial cells but rather prominent in stromal elements, including fibroblasts and endothelial cells (**Figure 2C**). Expression of SEMA3A receptor (*PLXNA1*) and co-receptor (*NRP1*) is rather promiscuous in PDAC tissues, which suggests that many cell types might be responsive to this axon guidance cue (**Figure S2A**). Next, we divided the samples of the PanCuRx cohort based on the *SEMA3A* expression status (either high or low, see methods) and further explored the association of *SEMA3A* with molecular features of aggressive PDAC. *SEMA3A*^high^ tumors were enriched for basal-like subtypes (**Figure S2B**) and major imbalances of the mutant *KRAS* allele (**Figure S2C**). We concluded that *SEMA3A* is linked to the basal-like/squamous subtype and that the different results observed for the TCGA cohorts (see Figure S1A) could be explained by the significant contamination from non-neoplastic cells. To corroborate that, we separated cases with the highest proportion of basal-like and classical epithelial cells from the scRNA-Seq of Peng et al. [40] (**Figure S1D**) to find that basal-like cells accordingly displayed elevated expression of *SEMA3A* (**Figure 2D**). Finally, we performed ISH analyses on human PDAC tissues (n=11) classified as either classical or basal-like/squamous based on the expression of markers of the two subtypes (**Figure S2E**). In keeping with the scRNA-Seq data (Figure 2C, Figure 2D), *SEMA3A* could only be detected in basal-like epithelial cells while detectable in stromal elements of both classical and basal-like tissues (**Figure 2E**). In sum, our analysis shows that *SEMA3A* expression is not restricted to epithelial cells in PDAC tissues, yet it is elevated in both stromal and epithelial compartments of basal-like tumors.

**Figure 2.**
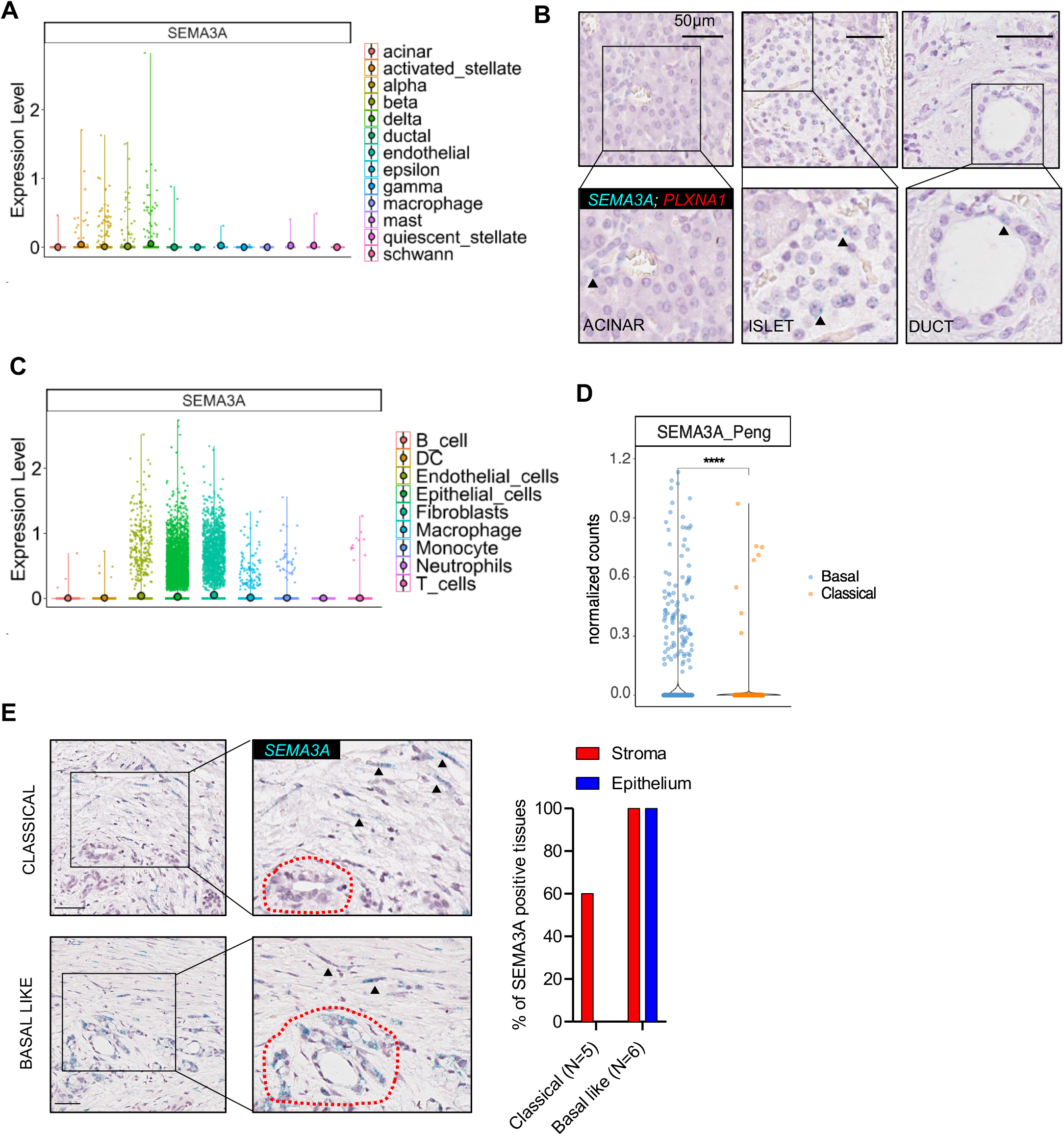
SEMA3A expression is selectively enriched in basal-like PDAC. **A**. Violin plots of the normalized expression of *SEMA3A* in each annotated cell cluster from the integration of 4 different scRNA-Seq datasets [35, 36, 37, 38] of normal pancreatic tissues (see methods). **B**. Representative ISH images showing expression of *SEMA3A* (green) and *PLXA1* (red) in acinar cells (left panel), islet cells (middle panel), and ductal cells (right panel). Scale Bar, 50 µm. Insets show magnification of selected areas, and arrowheads indicate epithelial cells with detectable expression of *SEMA3A*. **C**. Violin plots of the normalized expression of *SEMA3A* in each annotated cell cluster from the integration of 4 different scRNA-Seq datasets[8, 39, 40, 41] of pancreatic cancer tissues (see methods). **D**. Expression of *SEMA3A* in pancreatic ductal cells classified as classical or basal-like from Peng et al. [40]. See also Supplementary Figure 2D. ****, p < 0.0001 by Student t test. **E**. Representative ISH images showing expression of *SEMA3A* in pancreatic cancer tissues classified as classical (n = 5) or basal-like (n = 6). See also Supplementary Figure 2E. Right panel, bar plot displaying the percentage of cases that showed positivity for *SEMA3A* in the stroma (red) or epithelial (blue) compartment of the pancreatic cancer tissues.

### SEMA3A promotes anoikis resistance and a mesenchymal phenotype in mouse PDAC cells

To understand whether and how dysregulated SEMA3A levels contribute to promote malignancy of PDAC cells, we performed genetic perturbation experiments using both mouse PDAC cell lines and organoids. Since all the KPC derived cell lines expressed high levels of *Sema3a*, we first derived subclones displaying reduced expression of the gene (**Figure S3A**). To overexpress *Sema3a*, we stably transduced mT organoids and *Sema3a*^low^ subclones from both FC1199 and FC1245 (denoted by the B suffix) with a lentiviral vector carrying *Sema3a* open-reading frame (ORF) (**Figure 3A, 3B** and **Figure S3B-D**). Two different gRNA were instead transduced in inducible Cas9 expressing mM organoids as well as in parental FC1199 (designated by the A suffix). Cas9 was induced by treatment with doxycycline (2.5µg/µL) for 3 days. Given the link between *SEMA3A* expression and the TGF-β pathway, *Sema3a* mutant organoids were selected by withdrawal of TGF-β inhibitors (A83-01 and mNoggin) from the culture medium. Next, we verified knockout by measuring *Sema3a* expression at mRNA (Figure S3B, **Figure S3E)** and at protein level (Figure 3A, **Figure 3C**). Genetic manipulation of *Sema3a* resulted in coherent changes in the level of the secreted proteins in the cultures conditioned media (**Figure S3C**). Then, we evaluated the effect of different *Sema3a* levels on cell viability by measuring ATP cellular content over the course of 7 days of cultivation. We found no significant changes for both mouse organoids (**Figure 3D**) and monolayer cell cultures (**Figure 3E-F**). Next, we looked at the activation of relevant PDAC signaling pathways (i.e., MAPK and PI3K/Akt) by evaluating the phosphorylated levels of their main effectors. The overexpression of *Sema3a* in mT increased the activation of Akt, which was consistent with the reduced phosphorylation of Akt in mM organoids upon *Sema3a* knockout with two different gRNAs (**Figure 3G and Figure S3F and G**). The same pattern was observed in monolayer cell cultures, where elevated levels of SEMA3A were clearly associated with increased fluxes through the PI3K/Akt pathway (**Figure 3H, I**). More prominent in monolayer cell cultures than in organoids, the manipulation of *Sema3a* also affected fluxes through the MAPK pathway. Indeed, a significant reduction of phosphorylated levels of ERK1/2 could be observed upon depletion of *Sema3a* in FC1199A cells with two different gRNAs (**Figure 3I**). Considering the differences between the two culture systems, we can conclude that the dysregulation of *Sema3a* in mouse PDAC cells affects fluxes through the major signaling pathways. Given the correlation between SEMA3A expression and EMT gene programs in bulk transcriptomic datasets, we further explored whether overexpression/ silencing of *Sema3a* had causative role in defining a mesenchymal-like phenotype of PDAC cells. It is important to highlight intrinsic differences between cells grown as organoids or monolayer cell cultures, with the latter showing a prominent mesenchymal-like phenotype. Forced expression of *Sema3a* in mT resulted in increased levels of both *Snai1* and *Zeb1* at mRNA levels (**Figure S3H**) as well as of Vimentin at protein levels (**Figure 3G**). Less consistent effects on the expression of EMT genes/proteins were observed in mM following knockout of *Sema3a* (Figure 3G and Figure S3H). In mouse monolayer cell cultures, the effects of *Sema3a* dysregulation on the expression of EMT markers was overall negligible (**Figure 3H, 3I and Figure S3I, S3J**). Nonetheless, the manipulation of *Sema3a* expression in mouse PDAC cultures had significant effect on mesenchymal-like traits such as migration (**Figure 3J**) and anoikis resistance (**Figure 3K, 3L**). To evaluate the migratory capability of mouse PDAC cells of different genotypes, we performed the wound healing assay and found that *Sema3a* significantly promoted the migration of FC1199 cells (**Figure 3J**). In keeping with that, the treatment with recombinant SEMA3A rescued the effect of gene knockout on the migratory capacity of FC1199 cells (**Figure 3J right panel**). Next, we set up an anoikis assay for both monolayer cell cultures and organoids. Mouse metastatic organoids were detached from the extracellular matrix and then plated in PolyHEMA treated plates. Induction of apoptotic cell death was measured by evaluating Caspases 3/7 activation. The depletion of Sema3a from mM organoids significantly increased apoptotic cell death (**Figure 3K**). Exogenous supplementation of recombinant SEMA3A significantly reduced apoptotic cell death suggesting a prominent role for *Sema3a* in mediating this phenotype. In keeping with that, overexpression of *Sema3a* in FC1199 significantly reduced cell death of cells grown in suspension in PolyHEMA treated plates (**Figure 3L**). Although not statistically significant, the treatment with a selective Rho-associated kinase inhibitor (RhoKi, Y-27632)[42] seemed to partially counteract the beneficial effect of *Sema3a* overexpression on resistance to anoikis (**Figure 3L**). FAK is an important regulator of cell survival with an established role in mediating anoikis resistance [43]. Therefore, we measured the level of phosphorylation of the Tyrosine 397, the major autophosphorylation site of FAK, in cells displaying different levels of SEMA3A and/or treated with recombinant SEMA3A. We found that cancer-cell derived SEMA3A induces activation of FAK in both mouse organoids and monolayer cell cultures (**Figure 3M**). Accordingly, treatment of mouse PDAC cultures with recombinant SEMA3A stimulated FAK autophosphorylation (**Figure 3N**) and FAK inhibition with the selective inhibitor defactinib (FAKi) counteracted the SEMA3A protective effect against anoikis (**Figure 3O** and **Figure S3K**).

**Figure 3.**
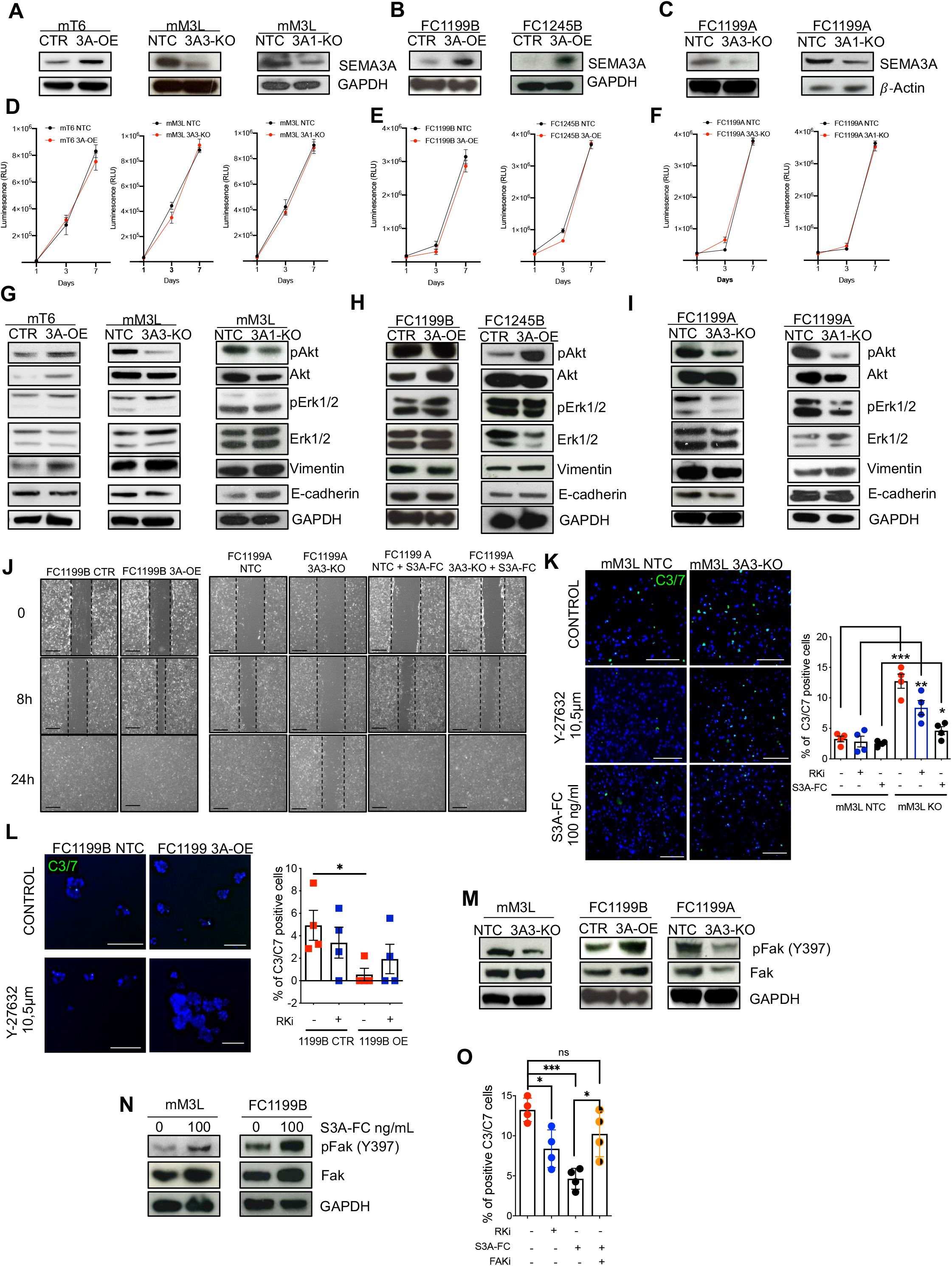
SEMA3A promotes anoikis resistance and a mesenchymal-like phenotype in mouse pancreatic cancer cells. **A-C**. Immunoblot analyses of SEMA3A in whole cell lysates from mouse tumor (mT) organoids, mouse metastatic (mM) organoids, and KPC 2D cell lines (FC1199 and FC1245) following either overexpression (OE) or genetic knockout (KO). GAPDH was used as loading control in A and B, while β-actin was used in C. The suffixes A and B for FC1199 and FC1245 denote clonal populations displaying high and low levels of SEMA3A, respectively. The suffix L for mM3 denotes that the culture was established from a liver metastasis. NTC or CTR denotes cultures stably transduced with a mock control. **D-F**. Proliferation (as total luminescence) measured over the course of 7 days of either cell lines or organoid cultures from A-C. **G-I**. Immunoblot analyses of the indicated proteins in whole cell lysates from mouse organoid cultures or mouse pancreatic cancer cell lines following either overexpression (OE) or genetic knockdown (KO) of SEMA3A. GAPDH was used as loading control. **J**. Representative photographs of the wound area taken immediately after (0), 8 and 24 hours after the incision for FC1199A and B cell lines stably transduced with either non-targeting or control vectors (NTC, CTR), SEMA3A ORF (OE), or gRNAs targeting SEMA3A (KO). FC1199A NTC and KO cells were also treated with recombinant SEMA3A. The experiment was performed in quadruplicate. **K**. Representative immunofluorescence images of the anoikis assay (see methods) performed on poly-HEMA coated plate for mM3L NTC and KO treated vehicle (Control), with a RhoK inhibitor (Y-27632), or with recombinant SEMA3A (S3A-FC). Scale bars, 100 µm. Quantification of 4 independent experiments is provided in the box plot on the right. ***, p < 0.001; **, p < 0.01; and *, p < 0.05 by Student t test. **L**. Representative immunofluorescence images of the anoikis assay (see methods) performed for FC1199B CTR and OE treated with vehicle (Control) or a RhoKi. Scale bars, 100 µm. Quantification of 4 independent experiments is provided in the box plot on the right. *, p< 0.05 by Student t test. **M**. Immunoblot analysis of phospho-FAK, and total FAK in whole cell lysates from mM cultures and KPC cell lines with different SEMA3A genotypes (e.g., KO or OE). **N**. Immunoblot analysis of phospho-FAK and total FAK in whole cell lysates from mM3L and FC1199B treated with recombinant SEMA3A. GAPDH was used as loading control in M and N. **O**. Quantification of apoptotic cells from the anokis assay of the SEMA3A knockout mM3L treated with vehicle, with the RhoK inhibitor (RKi), the recombinant SEMA3A (S3A-FC), or the combination of S3A-FC and defactinib (FAKi). Data are displayed as mean S.D. of 4 technical replicates. * p < 0.05; and ***. p < 0.001 by Student t test.

### SEMA3A expression sustains basal-like/squamous gene programs in PDAC

*In vitro* manipulation of *Sema3a* in mouse PDAC cells showed that the axon guidance cue promotes mesenchymal-like traits including enhanced migratory capacity and resistance to anoikis. To identify pathways downstream of *Sema3a* involved in promoting PDAC aggressiveness, we performed transcriptomic analysis on mouse PDAC cells of different genotypes. The overexpression of *Sema3a* had significant effects on the transcriptome of FC1199B with over 2,000 genes significantly up- and downregulated (**Figure 4A** and **Supplementary Table 1**). The genetic knock-out of *Sema3a* led to a similar degree of transcriptomic changes (**Figure 4D** and **Supplementary Table 3**). As shown in Figure 4A, the forced overexpression of *Sema3a* was associated with the significant downregulation of *Grem1*, a BMP inhibitor that has been shown to promote epithelialization of mesenchymal PDAC cells [44]. Next, we performed gene set enrichment analysis on the list of differentially expressed genes using the GSEA method [45] (**Figure 4B, 4E**). Following overexpression of *Sema3a*, we observed the enrichment of gene programs related to cytoskeleton remodeling and the activity of Rho GTPases (**Figure 4B, and Supplementary Table 2**). In keeping with that, the knock-out of *Sema3a* led to the reduced representation of the same gene programs (**Figure 4E, and Supplementary Table 4**). Secreted SEMA3A generally induces growth cone collapse in neurons by acting as either chemorepellent or chemoattractant through microtubule and actin reorganization [46]. Moreover, SEMA3A is reported to interact directly or indirectly with multiple GTPases, including Rho GTPases [46]. Of note, overexpression of *Sema3a* was also associated to a reduced representation of axon guidance gene sets, which we linked to the reduced expression of Slit/Robo genes. When looking at signatures of aggressive human PDAC, we found that SEMA3A^high^ cells presented higher “squamousness” than SEMA3A^low^ cells. Furthermore, SEMA3A deficient cell lines showed significant reduction of the GP3 gene programs defined by Bailey et al.[7] based on inferred activity of the TGF-β pathway (**Figures 4C-F**). Consistent with our findings in preclinical models, gene programs related to EMT, the TGF-β pathway, the activation of FAK, Rho GTPases, and wound healing were significantly enriched in *SEMA*^*hig*h^ tumors both in the ICGC [7] and the PanCuRx [8] cohorts (**Figure 4G-H, Supplementary Tables 5-8**).

**Figure 4.**
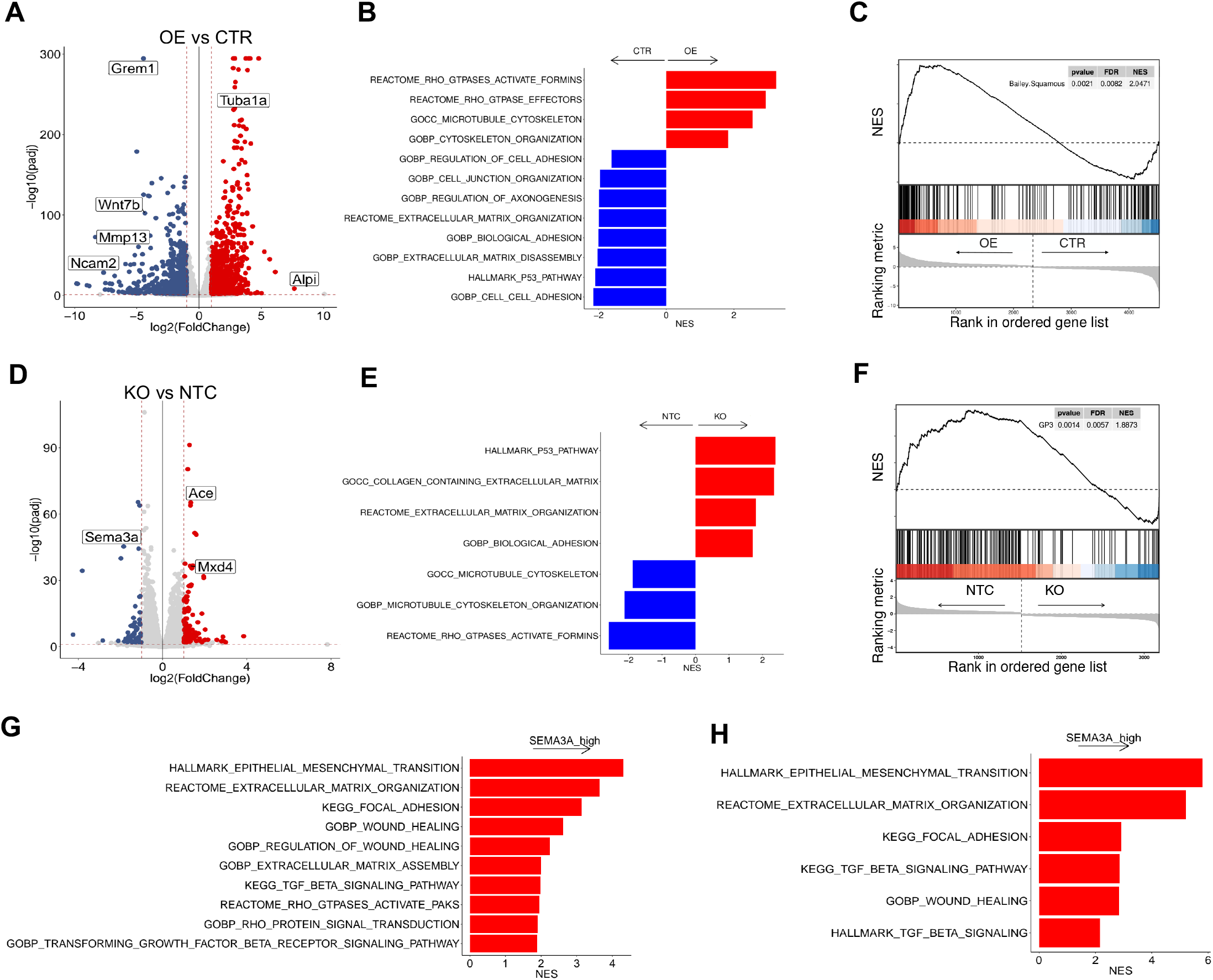
Transcriptomic changes following SEMA3A perturbation. **A**. Volcano plot of the differences in gene expression between control (CTR, n = 3) and *Sema3a* overexpression (OE n = 3). Indicated are some of the genes with log2FC expression ≥ 2 and adjusted *p* < 0.05. See Supplemental Table 1 for the full list of differentially expressed genes. **B**. Enrichment of selected pathways (GSEA) when comparing FC1199B *Sema3a*^*low*^ (CTR) and FC1199B overexpressing *Sema3a* (OE). See also Supplementary Table 2. **C**. GSEA plot evaluating the Squamous signature [7] upon *Sema3a* overexpression (OE) in FC1199B cells. **D**. Volcano plot of the differences in gene expression between control (NTC, n = 3) and *Sema3a* knockout (KO, n = 3). Indicated are some of the genes with log2FC expression ≥ 2 and adjusted *p* < 0.05. See Supplemental Table 3 for the full list of differentially expressed genes. **E**. Enrichment of selected pathways (GSEA) when comparing *Sema3a* proficient (NTC) and deficient (KO) FC1199A cells. See also Supplemental Table 4. **F**. GSEA plots evaluating the Gene Program 3 (GP3, TGFβ pathway)[7] upon *Sema3a* knockout in FC1199A cells. **G-H**. Enrichment of selected pathways when comparing SEMA3A high and low tissues from the ICGC[7] (G) and the PanCuRx[8] (H) cohorts. See also Supplementary Tables 5-8. In B, E, G, and H, GSEA was performed using gene sets from Hallmark, GO, KEGG, Reactome, and HP databases in MsigDB library. Displayed gene sets that passed false-discovery rate < 0.05.

### SEMA3A promotes PDAC progression *in vivo*

Under tissue culture conditions, we found that SEMA3A enhanced cell motility in scratch-wound assay and promoted resistance to anoikis (**Figure 3J, 3K**). Accordingly, integrated transcriptomic analyses showed that SEMA3A sustains EMT-related gene programs in PDAC. Next, we sought to assess the *in vivo* phenotypic consequences of *Sema3a* dysregulation. When injected into the tail vein of immunocompetent mice, *Sema3a* proficient cells rapidly colonized the lung parenchyma as opposed to *Sema3a* deficient cells (**Figure 5A**). Then, we evaluated tumor growth kinetics by transplanting *Sema3a* proficient and deficient cells into the pancreas of immunocompetent mice (n = 11/group) (**Figure 5B**). In terms of engraftment rate, two out of 11 mice transplanted with *Sema3a* deficient cells did not show any detectable mass while all the mice transplanted with *Sema3a* proficient cells eventually developed tumor masses (**Figure 5B**). Moreover, *Sema3a* expressing cells generated larger tumors following four weeks from transplantation (**Figure 5B**). To corroborate these findings, we decided to generate grafts based on the transplantation of mouse organoids which display delayed kinetics of *in vivo* tumor progression as opposed to monolayer cell cultures [28, 47]. In keeping with the transplantation of 2D cell lines, *Sema3a* deficient organoids generated smaller tumors in an immunocompetent host (**Figure 5C**). However, upon transplantation in immunodeficient hosts no difference was observed in the growth kinetics of *Sema3a* deficient and proficient cells (**Figure 5C**), which strongly suggest the involvement of the immune system in mediating the *in vivo* pro-tumorigenic effects of Sema3a. RNA-Seq analysis of tumor tissues collected at endpoint from mice transplanted with 2D cell lines did not reveal striking transcriptomic changes between the two groups (**Figure S4A and Supplementary Table 9**). However, gene-set enrichment analysis showed the overrepresentation of terms related to inflammation and interferon-related pathways in tumors lacking SEMA3A, which further pointed at cell extrinsic mechanisms (**Figure 5D and Supplementary Table 10**). Therefore, we generated additional cohorts of tumor-bearing mice using 2D cell lines and organoid cultures (mM3L) of different genotypes along with their controls. First, we measured the circulating levels of SEMA3A to confirm elevated blood levels of the protein in *Sema3a*-proficient tumor bearing mice (**Figure S4B, S4C**). Next, we performed immunohistochemistry for different immune markers and found that tumors generated by either cell lines or organoids lacking SEMA3A were characterized by reduced intra-tumor density of macrophages (F4/80+ cells) and increased infiltration by T cells (CD3+ and CD8+ T cells) (**Figure 5E, 5F**). In keeping with that, tumor tissues from immunocompetent mice transplanted with cell lines overexpressing SEMA3A were highly infiltrated by macrophages with reduced amount of intratumor T cells (**Figure S4D**). Finally, we analyzed the transcriptomic data from the ICGC cohort dividing tissues based on the expression of *SEMA3A* (see methods). Consistent with our findings in the mouse models, transcriptional signatures of different macrophages’ phenotypes were significantly enriched in *SEMA3A* high tumors (**Figure S4E)**.

**Figure 5.**
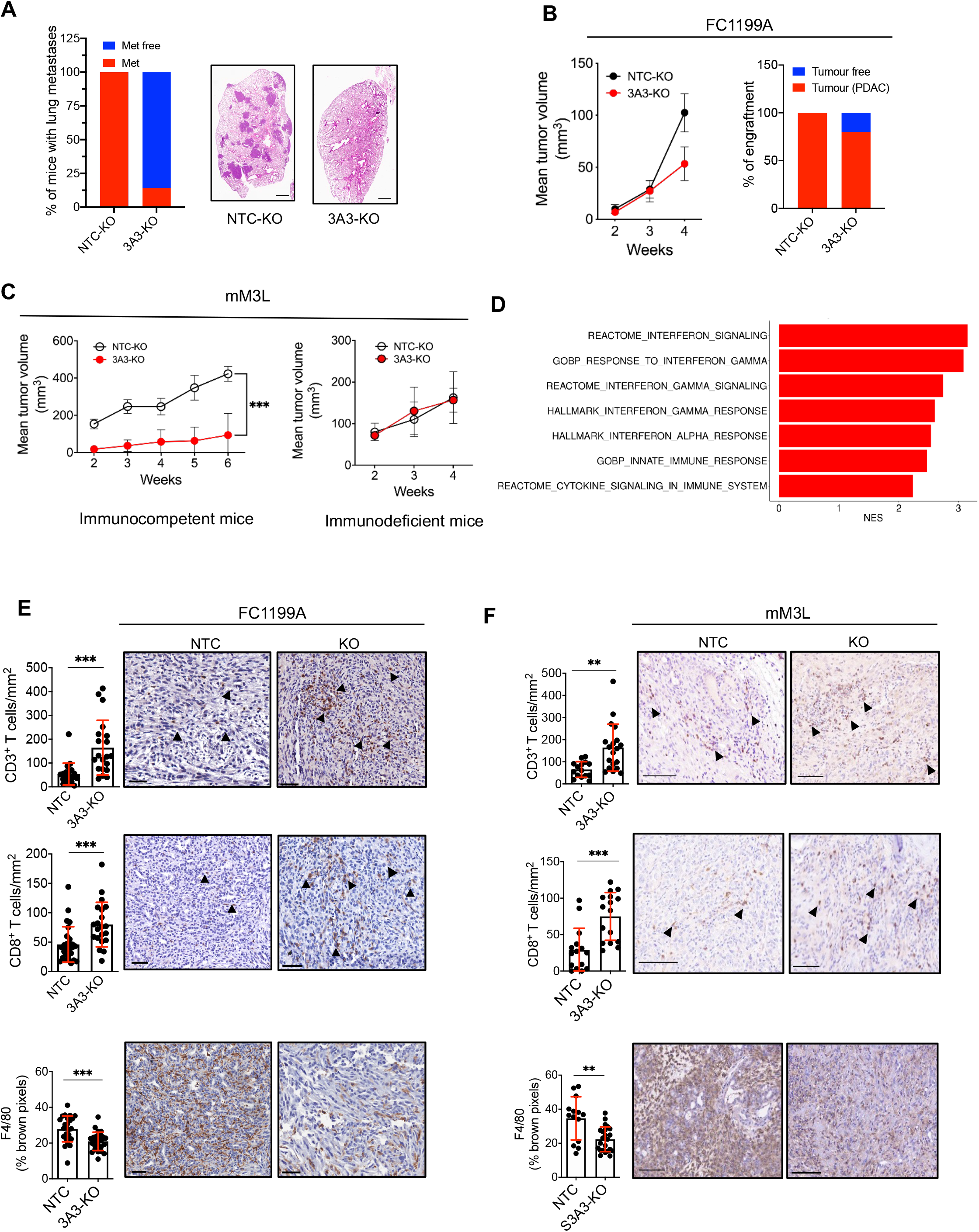
SEMA3A promotes growth of PDAC cells through modification of the tumor microenvironment. **A**. Stacked bar plot displaying the percentage of mice (n = 7 per group) displaying lung metastases upon tail-vein injection of *Sema3a* proficient (NTC) and deficient (KO) cells. Scale bar 1mm. **B**. On the left, line graph showing tumor volumes (mm^3^) of pancreatic masses detected upon the orthotopic injection of 1×10^6^ cells from FC1199A into immunocompetent mice (n = 11/group). Means ± S.D. are shown. On the right, stacked bar plot displaying the percentage of tumor bearing mice in the two cohorts (NTC and KO). **C**. Line graph showing tumor volumes (mm^3^) of pancreatic masses detected upon injection of 1×10^6^ cells from mM3L organoids into the pancreata of immunocompetent (n = 10 mice per group, left panel) or immunodeficient (n = 5 mice per group, right panel) mice. Means ± S.D. are shown. ****, p < 0.001 by two-way ANOVA with Sidak’s test for multiple comparison. **D**. Enrichment of selected pathways when comparing tissues from *Sema3a* deficient (n = 3) and proficient (n = 3) tumors. GSEA was performed using gene sets from Hallmark, GO, Reactome, and HP databases in MsigDB library. Displayed gene sets that passed false-discovery rate < 0.05. See Supplementary Tables 9 and 10 for details. **E-F**. Representative immunohistochemical staining for T cells markers (CD3 and CD8) and the macrophage marker F4/80 in pancreatic tissues from mice transplanted with: (E) FC1199A cells or (F) mM3L organoid cultures stably transduced with either non-targeting vector (NTC) or gRNA targeting *Sema3a* (KO). Scale bars, 50 µm. Quantification is provided on the left as mean ± S.D. (see methods). At least five individual areas per case and a minimum of five mice/arm were evaluated. Arrowheads indicate positive staining.

### Inhibition of CSF1R in SEMA3A high tumors maximizes chemotherapy benefit

Mouse PDAC tumors expressing high levels of SEMA3A are characterized by elevated infiltration of macrophages and reduced density of T cells. First, we investigated the effect of SEMA3A on macrophage’ activation and infiltration. Tumor-educated macrophages share features of M2-like macrophages, including high levels of Arginase 1. Conversely, M1-like macrophages are historically regarded as supportive of the antitumor immunity. To model the potential effect of SEMA3A on macrophages’ recruitment, we used the transwell migration assay. The monocyte/macrophage cell line RAW 264.7 was first polarized towards M1- or M2-like macrophages. Then, polarized and non-polarized macrophages were seeded with Matrigel in transwell to perform an invasion assay (see methods). As expected, a medium containing 20% FBS supported the invasion in all macrophages’ phenotypes (**Figure 6A**). Similarly, recombinant SEMA3A promoted the invasion of macrophages, particularly of M0 and M2-like phenotypes (**Figure 6A**). Then, we evaluated the effect of SEMA3A on the polarization of macrophages. Therefore, we grew RAW 264.7 in standard medium, in medium containing a cocktail of cytokine inducing the M2-like state (IL4 and IL13), and in conditioned media from SEMA3A proficient and deficient cells. As expected, the combined treatment with IL4 and IL13 induced morphological (**Figure 6B**) and molecular (**Figure 6C**) activation of the macrophages, which showed increased expression of *Arg1* and a slight (not significant) reduction of the expression of the M1-like gene *Nos2*. The conditioned medium from SEMA3A deficient and proficient cells induced the expression of *Arg1*, yet to a different extent. The exposure of cells to the conditioned medium from SEMA3A expressing cultures induced higher levels of *Arg1* as opposed to the conditioned medium from SEMA3A deficient cells, which instead also induced the M1-like gene *Nos2* (**Figure 6C**). Finally, the treatment of bone-marrow derived monocytes with recombinant SEMA3A induced protein and mRNA expression of M2-like markers (**Figure S5A, 5B**). In keeping with the *in vitro* data, we found a higher density of CD206+ macrophages in the tumor tissues established by *Sema3a* proficient cells (**Figure 6D**). Accordingly, tumor tissues from mouse PDAC cells overexpressing *Sema3a* displayed increased density of CD206+ macrophages (**Figure S5C**). To understand whether macrophages were responsible for the exclusion of T cells from the tumor bed of SEMA3A expressing cells, we depleted macrophages using a monoclonal antibody against CSF1R (αCSF1R). As shown in **Figure S5D**, immunocompetent mice were treated daily with αCSF1R three days prior the transplantation with SEMA3A proficient and deficient cells along with the control. The treatment with αCSF1R continued every other day until endpoint and tumor growth monitored by manual palpation and ultrasound imaging. At endpoint, we observed a significant reduction of intratumoral infiltration by macrophages (F4/80+ cells) in tumors from mice treated with αCSF1R regardless of the SEMA3A status (**Figure 6E, 6F**). Cytometric analyses of blood samples from tumor-bearing mice also confirmed the reduction of F4/80+ cells with no significant effect on Ly6C^+^Ly6G^+^ or Ly6C^+^ cells (**Figure S5E**), which is in line with the inhibition of CSF1R in mouse PDAC using a small molecule[6]. Only in tumors established by SEMA3A overexpressing cells, the depletion of macrophages was associated with increased intratumoral infiltration by CD8+ T cells (**Figure 6F**). These results suggest that in SEMA3A+ tumors, T cell exclusion is mainly mediated by macrophages. Next, we tested the effect of CSF1R inhibition alone or in combination with gemcitabine on the survival of mice bearing tumors from either *Sema3a* high or low cells (**Figure S5F**). In line with their less aggressive behavior, SEMA3A low tumors responded to all the treatments, yet gemcitabine monotherapy did not reach statistical significance (p = 0.08) (**Figure S5G**). SEMA3A high tumors responded poorly to both gemcitabine and CSF1R inhibition as monotherapy, and only the combination significantly extended the survival of the mice (**Figure 6G**).

**Figure 6.**
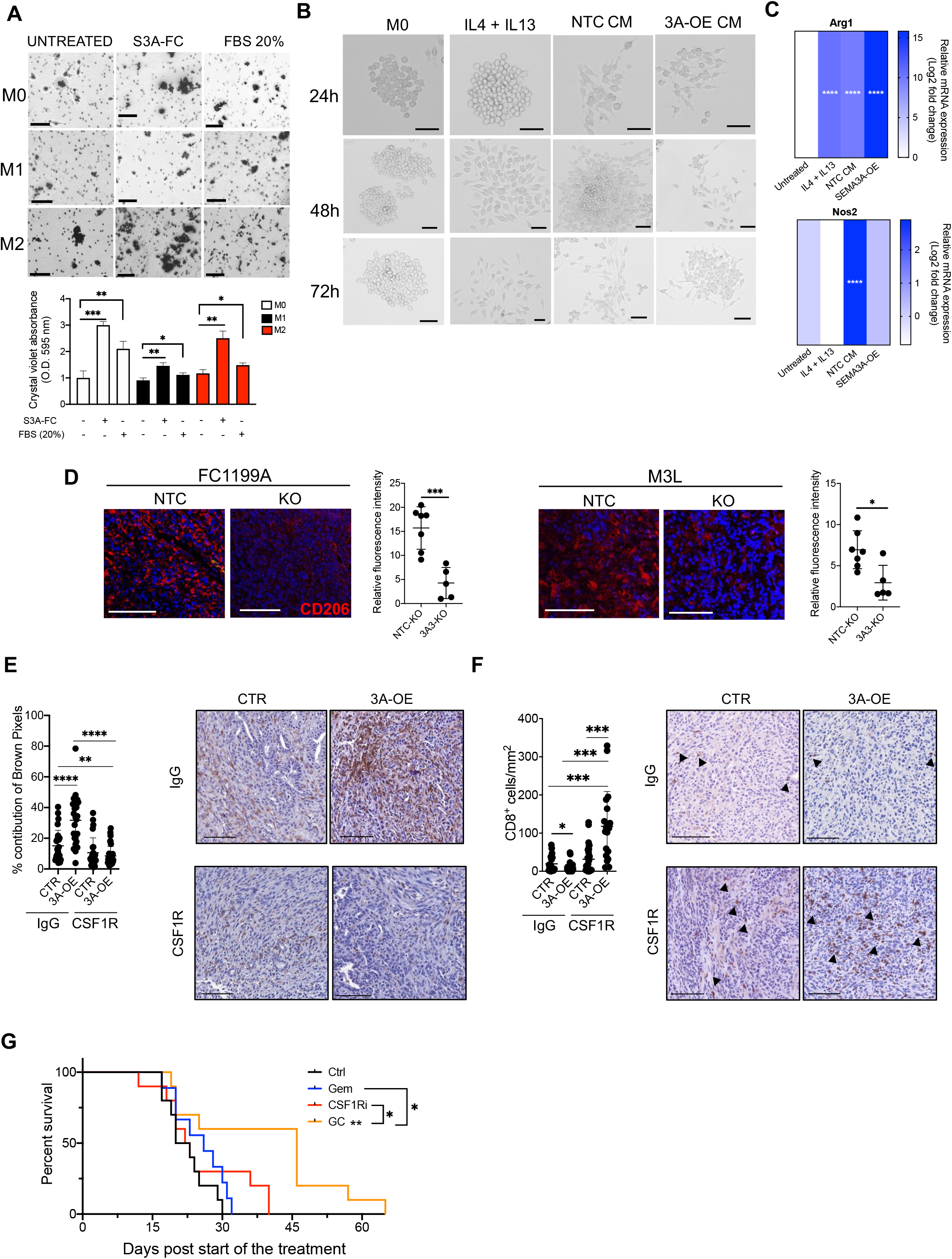
Targeting macrophages in SEMA3A expressing mouse tumors maximizes chemotherapy benefit. **A**. Bright-field images of migrated macrophages in the transwell assay (see methods). The quantification is provided at the bottom (bar plots) as mean ± S.D optical density values from three technical replicates. ***, p < 0.001; **, p < 0.01; and *, p < 0.05 by unpaired Student t test. **B**. Brightfield images of mouse RAW 264.7 cells treated with different conditions for up to 72 hours. From left to right, untreated (M0), combination of IL4+IL13, conditioned media from control cells, and conditioned media from cells overexpressing SEMA3A. **C**. Heatmap showing relative mRNA expression of *Arg1* (top) and *Nos2* (bottom) measure by qPCR of mouse cells from B. Data are mean of three technical replicates. **D**. Representative immunofluorescence staining for the M2-like marker CD206 (red) in tumor tissues from either FC1199 (left panels) or mM3L (right panels). Nuclei were stained with DAPI. Scale bars 100 µm. Quantification is provided on the right as relative fluorescence intensity (mean ± S.D.). At least three individual areas per case and a minimum of three mice/arm were evaluated. ***, p < 0.001; **, p < 0.01; and *, p < 0.05 by unpaired Student t test. **E**. Representative immunohistochemical staining for the macrophage marker F4/80 in pancreatic tissues from mice transplanted with FC1199B cells stably transduced with either mock (CTR) or a vector carrying *Sema3a* ORF (3A-OE) treated with control Ig G or CSF 1 R monoclonal antibody. **F**. Representative immunohistochemical staining for the cytotoxic T cell marker (CD8) in pancreatic tissues from mice transplanted with FC1199B cells stably transduced with either mock (CTR) or a vector carrying *Sema3a* ORF (3A-OE) treated with control IgG or CSF1R monoclonal antibody. In D and E, scale bars, 50 µm. Quantification is provided on the left as mean ± S.D. (see methods). At least five individual areas per case and a minimum of five mice/ arm were evaluated. Arrowheads indicate positive staining. **G**. Kaplan-Meier survival analysis of mice transplanted with SEMA3A high cells and treated with control IgG (Ctrl, n = 10), Gemcitabine (Gem, n = 10), αCSF1R (CSF1Ri, n = 10), or combination of Gemcitabine and αCSF1R (GC, n = 10). Statistical differences identified by log-rank test.

## Discussion

Genome-wide analyses of PDAC tissues have evidenced the dysregulation of the axon guidance signaling pathway in this dismal disease [12, 14, 48, 49]. Here, we investigated the role of the diffusible axon guidance cue SEMA3A, whose tissue expression has been previously found to be elevated in patients with poor clinical outcome [12, 13]. We show that SEMA3A is highly expressed by neoplastic cells with squamous differentiation and a basal-like phenotype. Of the two PDAC cell lineages [7, 8, 9, 10], the basal-like/squamous phenotype displays a more aggressive behavior and it is enriched in post-treatment tumors as well as in metastases [8]. We found that both cell-intrinsic and cell extrinsic factors promoting the basal-like/squamous subtype induce expression of *SEMA3A* in PDAC cells. Mechanistically, we demonstrate that SEMA3A exerts both cell-autonomous and non-cell autonomous effects to support the progression of PDAC. Cell-intrinsically, SEMA3A activates FAK and gene programs related to cytoskeleton remodeling to promote anoikis resistance and the migratory phenotype of PDAC cells. In keeping with that, SEMA3A overexpressing mouse PDAC cells display superior metastatic competence compared to cells lacking SEMA3A. Moreover, SEMA3A expressing cells induces pro-tumorigenic changes in the tumor microenvironment with increased density of macrophages and significantly reduced infiltration of T cells. Tumor-associated macrophages are the most abundant leukocyte population in the stroma of both mouse and human PDAC [6] and they contribute to establish an “immunologically cold” microenvironment also through T cell exclusion. Specifically in the context of SEMA3A expressing tumors, the depletion of macrophages led to increased intratumoral infiltration of T cells and the maximization of therapeutic benefit from gemcitabine.

The axon guidance is a highly conserved pathway involved in the proper formation of neural circuits during the development of the central nervous system (CNS)[46]. Furthermore, the activity of this pathway contributes to the maintenance and plasticity of neural circuits throughout the lifetime. The axon guidance genes include membrane-bound or diffusible ligands (Netrins, Semaphorins, Ephrins, Slits) that act either as chemoattractant or chemorepellent for growing axons and migrating neurons. These axon guidance cues and their receptors are also expressed outside of the CNS where they regulate cell-to-cell, cell-to-extracellular matrix interactions, and tissue morphogenesis [50]. At molecular level, all guidance cues influence cell motility through the engagement of the Rac family of small GTPases [50]. Here, we found that SEMA3A sustains gene programs related to EMT, cytoskeleton remodeling, and Rho GTPase signaling in mouse PDAC cells. The activation of those gene programs parallels a functional phenotype of mesenchymal-like cells with migratory capability and increased metastatic competence. Most of previous studies in PDAC has focused on investigating the role of members of the Slit/Robo axis on the PDAC malignant traits of PDAC as well as its cell identity [15, 18, 19, 20, 21]. According to their role in maintaining pancreatic cell identity [18], members of the Slit/Robo pathway have been reported to exert *bona fide* tumor suppressor mechanisms that limits stromal activation in disease [20]. However, other members of the pathway (i.e., ROBO3) have been shown to promotes the basal-like phenotype through a non-canonical signaling pathway [15]. Of the four classes of ligands, Semaphorins represent the largest family and were originally identified as chemorepellent proteins in the nervous system [22, 23]. Semaphorins can be secreted, GPI-anchored, or transmembrane proteins with recognized functions outside the CNS, including the regulation of angiogenesis and immune responses [24] [25]. SEMA3A belongs to the Class 3 of secreted Semaphorins and its potential role in cancer still needs to be elucidated. Indeed, several works have proposed a tumor suppressive role for SEMA3A, which has been reported to restrain tumor growth by hampering tumor angiogenesis [51]. In PDAC, an NRP1-independent superagonist SEMA3A was used as vasculature normalizing agent which demonstrated anti-tumor activity (Gioelli et al., 2018). Moreover, there are contradictory results on the effect of SEMA3A on recruitment and activation of tumor-associated macrophages (TAMs). TAMs have an established pro-tumoral function and shares features with M2-like macrophages, including the expression of Arginase 1 and of the Mannose Receptor CD206 [52, 53]. Carrer et al. reported that SEMA3A recruits a subset of resident Nrp1+ anti-tumoral macrophages [54], while Casazza et al. found that SEMA3A entraps protumoral macrophages in highly hypoxic areas [55]. Finally, Wallerius et al. reported a differential effect of SEMA3A on the proliferation of M2 and M1-like macrophages [56]: SEMA3A favored the expansion of anti-tumoral M1-like macrophages which was associated with the recruitment of cytotoxic T cells and a tumor-inhibiting effect. In our preclinical models, SEMA3A expressing cells increased the intratumoral density of macrophages while reducing infiltration by CD8+ T cells. Our findings perfectly align with the elevated expression of *SEMA3A* in basal-like/squamous PDAC, which are characterized by elevated infiltration of TAM and scant T cells [6]. *In vitro*, SEMA3A functioned as chemoattractant for different macrophages subsets and further skewed the macrophage population towards an M2-like phenotype. Accordingly, the depletion of macrophages with the monoclonal antibody against CSF1R favored intratumoral infiltration of cytotoxic T cells specifically in the context of SEMA3A^high^ tumors. Furthermore, we found that SEMA3A^high^ tumors were more resistant to gemcitabine treatment than the SEMA3A^low^ tumors, which perfectly aligns with the more aggressive biological behavior of SEMA3A^high^ tumors. However, the depletion of macrophages resulted in a significant greater benefit in terms of overall survival following chemotherapy for tumors with high expression of SEMA3A. Overall, we show here that SEMA3A is a functional marker of aggressive PDAC that promotes tumor progression by enhancing the metastatic competence of neoplastic cells and the macrophage-mediated exclusion of cytotoxic T cells. However, a greater infiltration of CD8+ T cells is observed in SEMA3A^high^ tumors upon macrophages depletion, suggesting a potential chemoattractant role of SEMA3A for T cells. While this aspect needs further elucidation, we have provided proof that SEMA3A^high^ tumors might benefit from therapies combining targeting of macrophages and chemotherapy.

## Supporting information

Supplementary file description

## Acknowledgment

We gratefully acknowledge the Centro Piattaforme Tecnologiche (CPT - University of Verona, Verona, Italy) for granting access to the genomic facility of the University of Verona. Additionally, we are grateful to Sonia Grimaldi, Nicola Sperandio, and Giada Bonizzato (ARC-Net Research Centre, University of Verona) for the assistance provided with the generation of organoid models.

## Authors’ Contribution

FL, SU, and VC conceived and designed the research; FL and VC developed the idea; FL, EF, and MB performed animal experiments; FL, and LV performed experiments with primary CAFs culture and organoids; FL and CH performed in vitro experiments; AA, CF, and FdS performed cytofluorimetric analysis; FL, AA, CF performed experiments with monocytes and macrophages; FL and AM performed ISH experiments; PD, PB, FP, DP analysed omics data and generated displays; SDa and SA established human organoids cultures; RTL, CC, and GP collected tissue samples. FL, EF, and LV performed histology and immunohistochemistry of all human and mouse tissue samples. AS, FL, and VC analysed the human tissues; IA assisted with the design of Chip-qPCR experiments and the interpretation of the data; FL, SU, FdS, and VC analysed data relative to the mouse TME; DT provided human and mouse models; SU and VC designed the in vivo treatment experiments; FL, SU, and VC interpreted the data; FL, FP, and VC wrote the manuscript. VC supervised the study.

