## Supplementary figures and images for "The axon guidance cue SEMA3A promotes the aggressive phenotype of basal-like PDAC"

### Supplementary figures

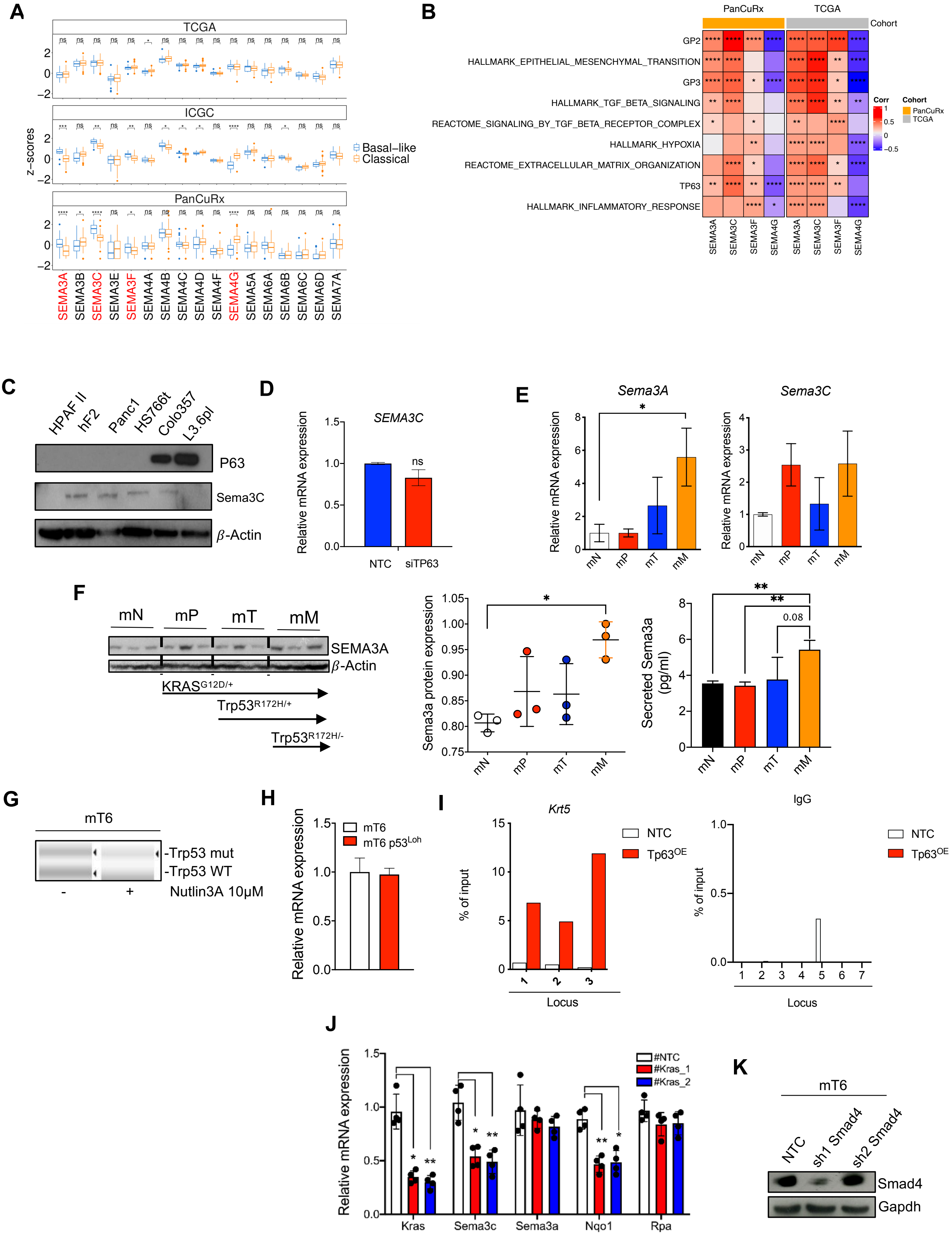

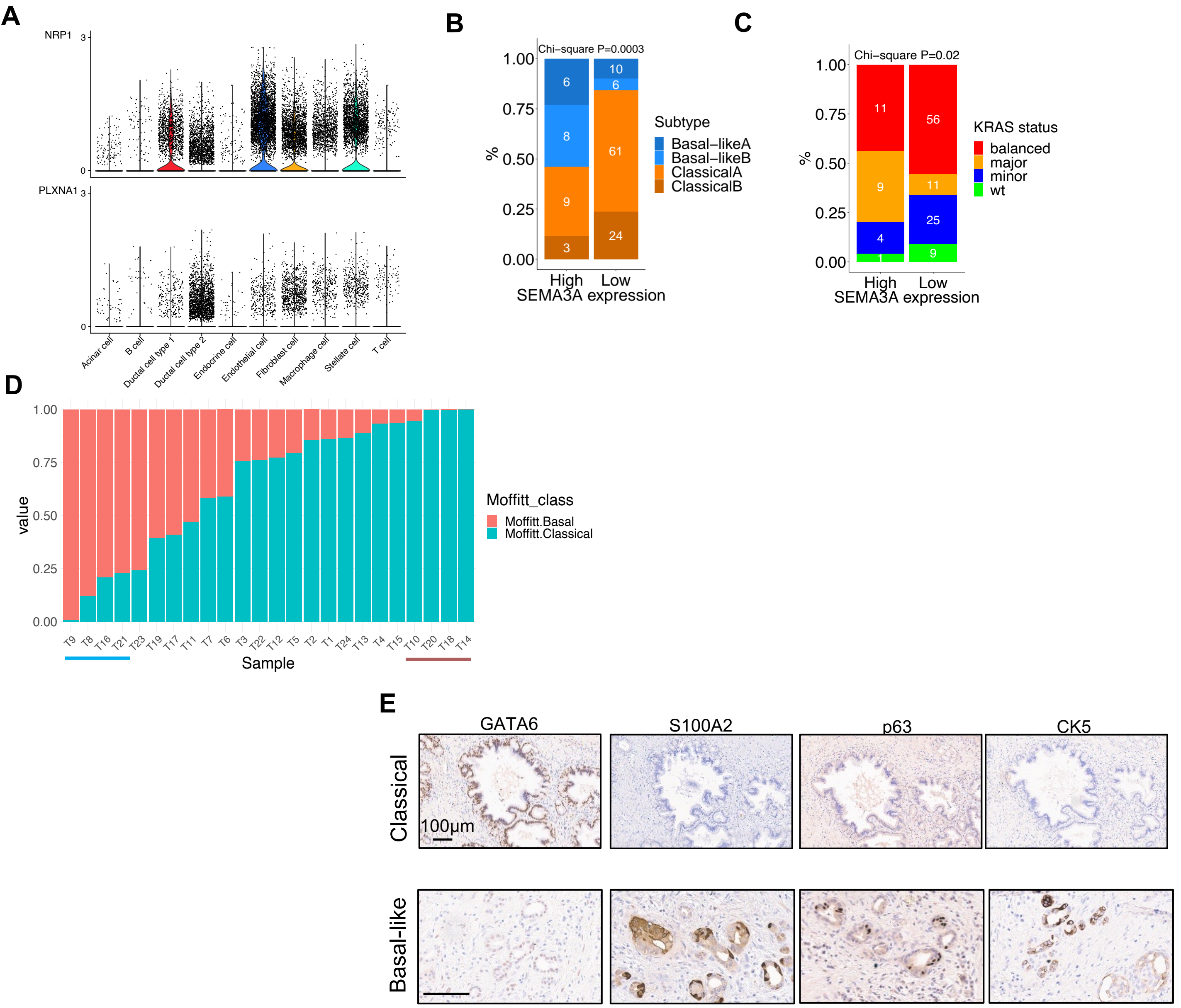

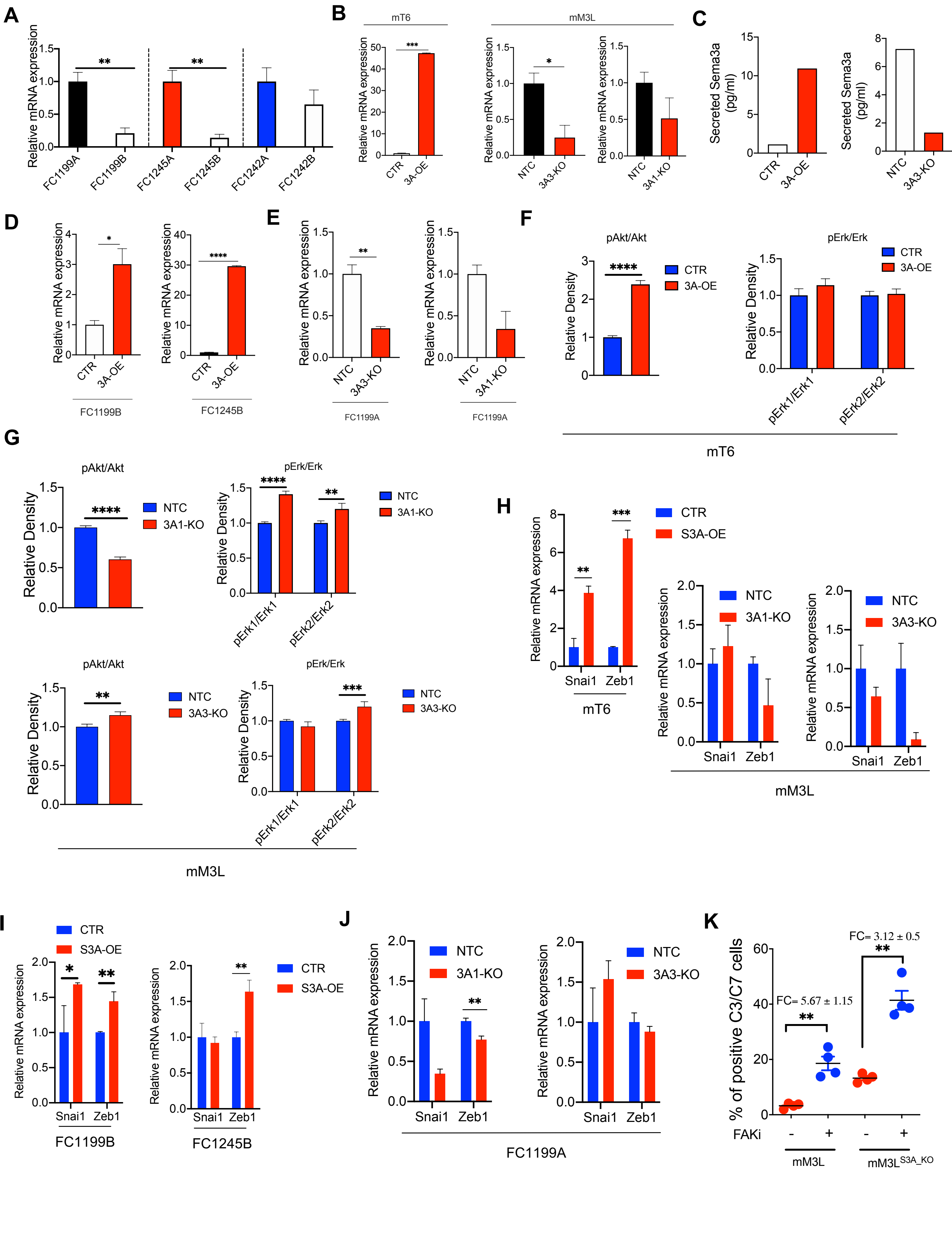

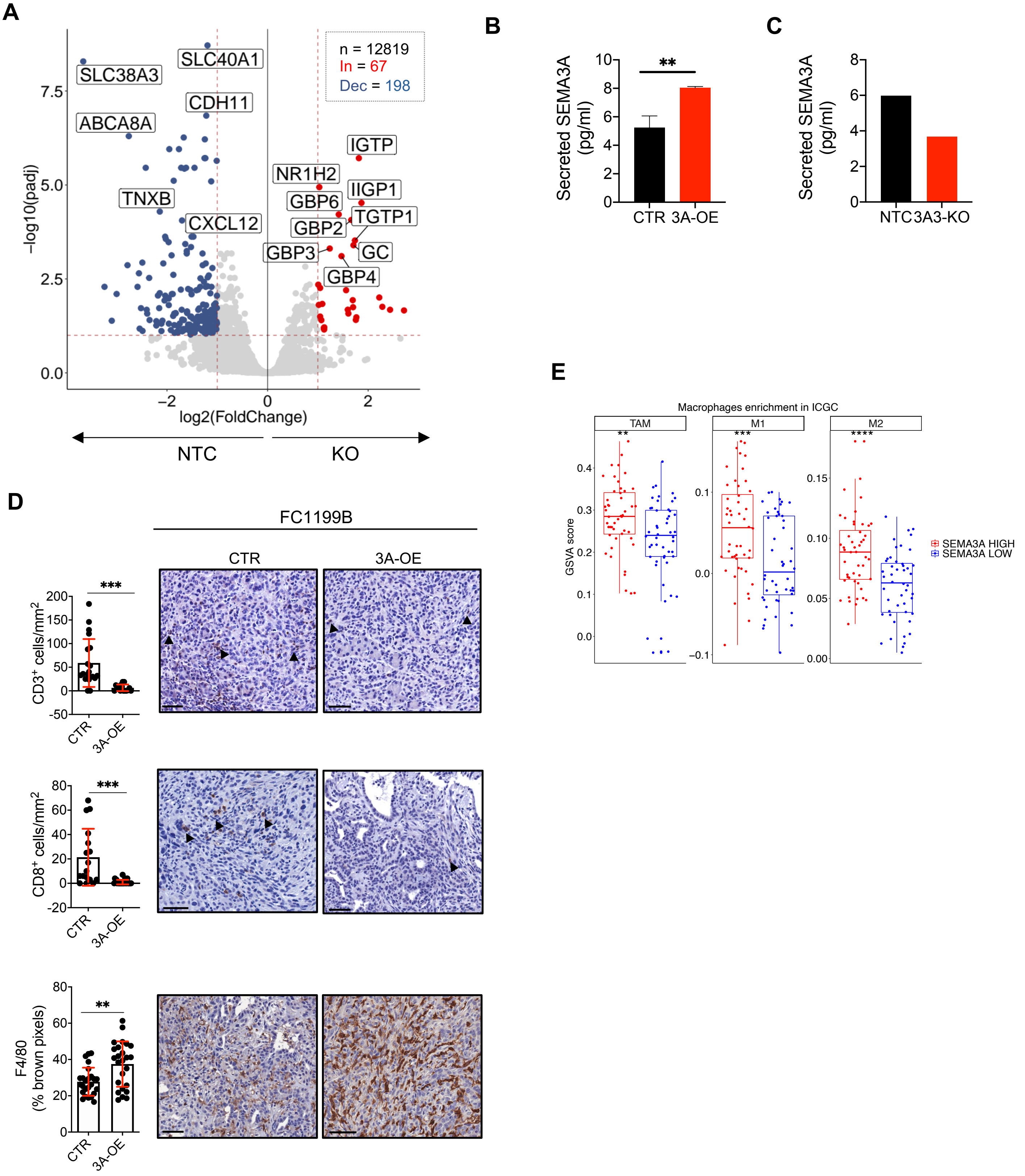

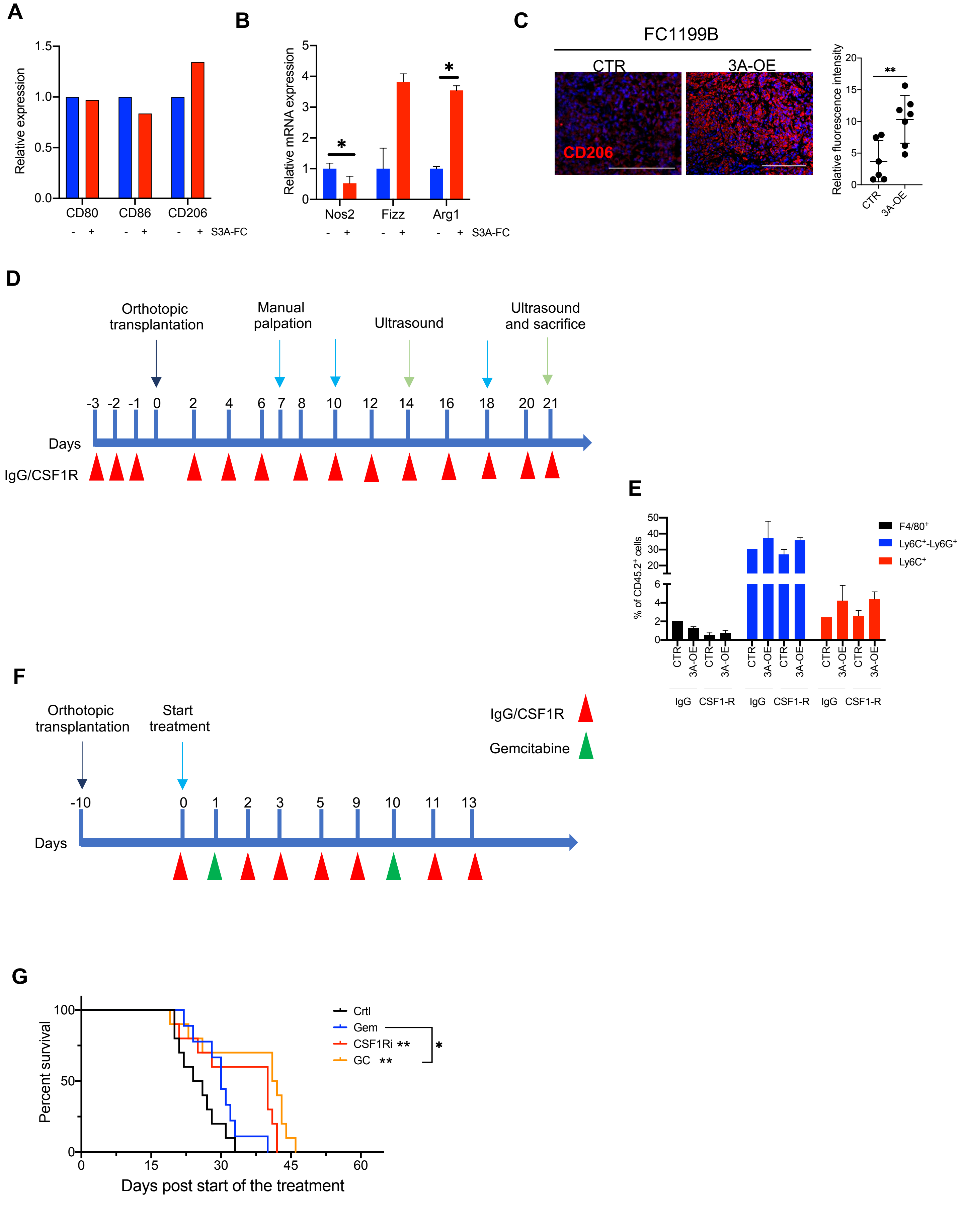
