## Supplementary file description for "The axon guidance cue SEMA3A promotes the aggressive phenotype of basal-like PDAC"

**SUPPLEMENTAL TABLES DESCRIPTION**

**Table S1.** List of differentially expressed genes upon *Sema3a* overexpression in FC1199 cells.

**Table S2**. Gene-set enrichment analysis (GSEA) on the list of differentially expressed genes upon Sema3a overexpression in FC1199 cells. Terms highlighted in yellow are reported in the relative figures.

**Table S3**. List of differentially expressed genes upon *Sema3a* knock-out in FC1199 cells.

**Table S4**. Gene-set enrichment analysis (GSEA) on the list of differentially expressed genes upon Sema3a knockout in FC1199 cells. Terms highlighted in yellow are reported in the relative figures.

**Table S5.** List of differentially expressed genes from the comparison between SEMA3A high and low tumors of the ICGC cohort.

**Table S6**. GSEA analysis on the list of differentially expressed genes from Table S5. Terms highlighted in yellow are reported in the relative figures.

**Table S7**. List of differentially expressed genes from the comparison between SEMA3A high and low tumors of the PanCuRx cohort.

**Table S8**. GSEA analysis on the list of differentially expressed genes from Table S7. Terms highlighted in yellow are reported in the relative figures.

**Table S9**. List of differentially expressed genes from mRNA-seq analysis of tissues from Sema3a proficient and deficient tumors.

**Table S10**. Gene-set enrichment analysis (GSEA) on the list of differentially expressed genes from Table S9. Terms highlighted in yellow are reported in the relative figures.

**Table S11.** Custom Macrophages' gene expression signatures.

### SUPPLEMENTAL FIGURE LEGENDS

**Supplementary Figure 1. Regulation of *Sema3a* and *Sema3c* expression by cell intrinsic and cell extrinsic inputs. A.** Boxplot of Semaphorins Z-scores stratified by the Moffitt subtypes [1] in the TCGA[2], the ICGC[3], and the PanCuRx[4] cohorts. *, $p < 0.05$; **, $p < 0.01$; and ***, $p < 0.001$ by Student t test. Only Semaphorins whose expression could be detected in all the transcriptomic datasets are shown; the Semaphorin genes displayed in Figure 1A are in red colored font. **B**. Heatmap showing correlation (Spearman's correlation) between the indicated Semaphorins and basal-like/ squamous associated gene programs in the PanCuRx[4] and the TCGA[2] cohorts. All annotated boxes, $p < 0.05$. **C.** Immunoblot analysis of p63 and SEMA3C expression in whole-cell lysates from established human cell lines. β-actin was used as loading control. **D**. Changes in the expression of *SEMA3C* in L3.6pl cells following knockdown of p63. **E**. qPCR analysis of *Sema3a* and *Sema3c* expression in mN (n = 3), mP (n=3), mT (n=6), and mM (n = 3) organoids. Results are shown as mean ± SD. *, $p < 0.05$ as determined by Student's *t*-test. **F.** Immunoblot analysis of SEMA3A in whole cell lysates from mN, mP, mT, and mM organoids. β-actin was used as loading control (left panel). Reported below the immunoblot, the *Kras* and *Trp53* status of the cultures. Middle panel, scatter dot plot showing the quantification of SEMA3A from the immunoblot on the right. *, $p < 0.05$ by Student t test. Right panel, levels of SEMA3A detected in the secretome of mouse organoid cultures using an ELISA assay. **, $p < 0.01$; *, $p < 0.05$; and 0.08 by Student t test. **G.** Snapshot of the gel-like image output of the DNA-chip bioanalyzer run showing the loss of the *Trp53* wild-type allele upon Nutlin-3A treatment (see methods) of mT organoids. Arrowheads indicate the bands corresponding to either wild-type or mutant Trp53. **H**. qPCR analysis of *Sema3c* in mouse tumor organoids displaying loss-of-heterozygosity of p53 compared to the parental culture. **I**. Anti-p63 ChIP-qPCR analysis of 3 different genomic regions of the *Krt5* promoter from mouse PDAC cells transduced with either empty control (NTC) or p63 ORF. On the right, anti-IgG Chip-qPCR of the 7 genomic regions of Figure 1G. **J**. Changes in the expression levels of *Kras, Sema3c, Sema3a, Nqo1,* and *Rpa3* in mouse PDAC cell lines transfected with empty vector or two different siRNAs targeting the mutant Kras allele. Results are shown as mean ± S.D. of 4 independent experiments. **, $p < 0.01$; and *, $p < 0.05$ by Student t test. **K**. Immunoblot analysis of Smad4 in mouse tumor organoids transduced with empty vector or with shRNAs (#1 and #2) targeting SMAD4. GAPDH was used as loading control.

**Supplementary Figure 2. Expression of SEMA3A in PDAC tissues. A**. Violin plots of normalized expression of *NRP1* and *PLXNA1* in each annotated cell cluster from Peng et al. [5]. Cells annotated as ductal cell type 2 correspond to neoplastic cells. **B**. Stacked bar plot showing the molecular subtypes distribution according to the *SEMA3A* expression status (see methods) in the PanCuRx [4] cohort. **C**. Stacked bar plots showing the distribution of *KRAS* genetic status according to the expression of *SEMA3A* in the PanCuRx [4] dataset. **D**. Samples from Peng et al. [5] ranked from left to right based on the proportion of basal-like or classical cells. For the analysis in Figure 2D, we considered the cells from the cases in blue (n = 4) and red (n = 4) which show the highest proportion of basal-like and classical cells, respectively. **E**. Representative immunohistochemical staining of classical (GATA6) or basal-like (S100A2, TP63, CK5) markers in pancreatic cancer tissues used to assess stromal and epithelial expression of *SEMA3A* in Figure 2E. Scale bars, 100 µm.

**Supplementary Figure 3. SEMA3A promotes the EMT phenotype in mouse pancreatic cancer cells. A.** qPCR analysis of *Sema3a* in parental KPC cell lines and derived subclones. **, p < 0.01 by Student t test. **B.** qPCR analysis of *Sema3a* in mT6 after transduction with a vector carrying *Sema3a* ORF (OE, left panel), and in mM3L following stable transduction with two different gRNAs targeting *Sema3a* (KO, right panels). ***, p < 0.001; *, p < 0.05 by Student t test. **C**. Levels of secreted SEMA3A assayed by ELISA in the conditioned media from mT6 (CTR and OE) and mM3L (NTC and KO). **D**. qPCR analysis of *Sema3a* in two different KPC cell lines following transduction with a vector carrying the *Sema3a* ORF. ***, p < 0.001; *, p < 0.05 by Student t test. **E**. qPCR analysis of *Sema3a* in FC1199A cell line following knockout with two different gRNAs targeting *Sema3a.* **, p < 0.01 by Student t test. **F**. Bar plots showing the quantification of changes in the phosphorylated levels of p-AKT (left panel) and p-ERK1/2 (right panel) as relative density of the total protein level. Data presented as means ± SD of three biological replicates. ****, *p* < 0.001 by Student's *t*-test corrected for multiple comparison using the Holm-Sidak method. **G**. Bar plots showing the quantification of the changes in the phosphorylated levels of p-AKT and p-ERK1/2 (relative density of the total protein) in mM3L following knockout with two different gRNAs. In H and I, **, p < 0.01; ***, p < 0.001; ****, p < 0.0001 by Student's *t*-test corrected for multiple comparison using the Holm-Sidak method. **H**. Changes in the expression levels of EMT genes (*Snai1*, *Zeb1*) in mT6 and mM3L organoid cultures following either overexpression or knockout of *Sema3a*. **I-J**. Changes in the expression level of EMT genes (*Snai1, Zeb1*) in FC1199 and FC1245 following either overexpression or knockout of *Sema3a*. ***, p < 0.001; **, p < 0.01, and *, p < 0.05 by Student t test (in E to G). **K**. Quantification of the anoikis assay (% of cleaved Caspase3/7 positive cells) of the mM3L wild-type and knockout for *Sema3a*
in the presence or the absence of the FAK inhibitor defactinib. Indicated the fold change (FC)± S.D. between treated/untreated for each genotype. **, p < 0.01 by Student t test.

**Supplementary Figure 4. *In vivo* effect of SEMA3A perturbation. A**. Volcano plot of the differences in gene expression between control (NTC, n = 3) and *Sema3a* knockout models (KO, n = 3). Indicated are some of the genes with log2FC expression ≥ 2 and adjusted *p* < 0.05. Provided in the box is the number of totals, upregulated (red), and downregulated (blue) genes. See Supplemental Table 9 for the full list of differentially expressed genes. **B**. Levels of circulating SEMA3A detected in the blood of mice transplanted with FC1199B stably transduced with either empty (CTR) vector or *Sema3a* ORF (OE). Results are shown as mean ± S.D. of three biological replicates. **, p < 0.01 by unpaired Student t test. **C**. Levels of circulating SEMA3A detected in the blood of mice transplanted with FC1199A stably transduced with a non-targeting control vector (NTC) or gRNA targeting *Sema3a* (KO). Results are shown as mean of two biological replicates. **D.** Representative immunohistochemical staining for CD3, CD8, and F4/80 of pancreatic tissues from mice transplanted with FC1199A stably transduced with either mock (CTR) or a vector carrying *Sema3a* ORF (OE). Scale bars, 50 µm. Quantification is provided on the left as mean ± S.D. (see methods). At least five individual areas per case and a minimum of five mice/arm were evaluated. **F**. Boxplot of macrophages related signature scores in tumors from the ICGC cohort[3] according to the expression of SEMA3A. p-values, Wilcoxon rank-sum test. ****, p < 0.0001; ***, p < 0.001; and **, p < 0.01. See also Supplementary Table 11.

**Supplementary Figure 5. CSF1R inhibition in mouse PDAC tumors. A.** qPCR analysis of *CD80*, *CD86* and *CD206* in bone-marrow derived monocytes treated with vehicle (-) or recombinant SEMA3A (+) (100 ng/mL) for 72 hours. Experiment was performed twice, and data are shown as mean value relative the untreated cells. **B** qPCR analysis of *Nos2*, *Fizz*, *Arg1* in bone-marrow derived monocytes treated with vehicle (-) or recombinant SEMA3A (+) (100 ng/mL) for 72 hours. Data are shown as mean ± S.D of three independent experiments. Statistical significance by Student t test: *, p < 0.05. **C**. Representative immunofluorescence staining for CD206 in tissues from tumor-bearing mice transplanted with FC1199 transduced either with control vector (CTR) or *Sema3a* ORF (3A-OE). Scale bars 100 µm. Quantification is provided on the right as relative fluorescence intensity (mean ± S.D.). At least three individual areas per case and a minimum of three mice/arm were evaluated. **, p < 0.01 by unpaired Student t test. **D**. Experimental design for the evaluation of CSF1R monoclonal antibody in a mouse model of PDAC. **E**. The effect of CSF1R inhibition on the depletion of monocytes in the blood circulation. *, p < 0.01 by Student t test. **F**. Experimental Design for the survival study of tumor-bearing mice treated with vehicle (IgG), CSF1R monoclonal antibody, Gemcitabine, or combination of CSF1Ri and Gem. **G**. Kaplan-Meier survival analysis of mice transplanted with SEMA3A low cells and treated with control IgG (Ctrl, n = 10), Gemcitabine (Gem, n = 10), αCSF1R (CSF1Ri, n = 10), or combination of Gemcitabine and αCSF1R (GC, n = 10). Statistical differences identified by log-rank test.

### MATERIALS AND METHODS

#### Human Samples

Fresh and formalin-fixed and paraffin-embedded (FFPE) PDAC tissues used in this study were obtained from surgical resection of patients treated at the University and Hospital Trust of Verona (Azienda Ospedaliera Universitaria Integrata, AOUI). All tissue specimens were acquired from treatment-naive patients and written informed consent from the donors for research use of the tissue was obtained prior to acquisition of the specimens. FFPE tissue of 11 individual PDAC cases was retrieved from the ARC-Net Biobank and were collected under the protocol number 1885 approved by the local Ethics Committee (*Comitato Etico Azienda Ospedaliera Universitaria Integrata*) to A.S (Prot. 52070, Prog. 1885 on 17/11/2010). These specimens were used for *in situ* hybridization and immunohistochemical analyses. Fresh tissue from curative resections was used for the generation of organoids and was collected under the protocol number 1911 approved by the local Ethics Committee (Comitato Etico Azienda Ospedaliera Universitaria Integrata) to V.C. (Prot. n 61413, Prog 1911 on 19/09/2018). All experiments were conducted in accordance with relevant guidelines and regulations.

#### Cell lines, organoids, and culture conditions

A total of 10 human pancreatic cancer cell lines were used in this study. The human PDAC cell lines HPAF-II, PANC1, and AsPC-1 were purchased from ATCC (CRL-1997, CRL-1469, CRL-1682). SUIT-2, Hs766T, L3.6pL, COLO 357, and BxPC3 cell lines were kindly provided by Prof. Aldo Scarpa (University of Verona). hF2-2D (monolayer) was kindly provided by Dr. David A. Tuveson (Cold Spring Harbor Laboratory, USA). HPAF-II, SUIT-2, PANC-1, L3.6PL, COLO 357, and BxPC3 were grown in DMEM (Gibco) supplemented with 10% FBS and 1% Penicillin-Streptomycin (Pen-Strep, Gibco). Hs766T, AsPC-1, and hF2-2D were cultured in RPMI (Gibco) supplemented with 10% FBS and 1% Pen-Strep. The two mouse PDAC cell lines used herein (FC1199 and FC1245) were established from tumor tissues of KPC (*Kras*$^{LSL\text{-}G12D/+}$;*Trp53*$^{LSL\text{-}R172H/+}$;*Pdx-1*-Cre) mice [6] and grown in DMEM supplemented with 10% FBS and 1% Penicillin-Streptomycin (Pen-Strep, Gibco). Human (n = 5) and mouse (n = 12) pancreatic organoids were established as previously described [7, 8]. For certain experiments, organoids were adapted to grow either on plastic or embedded in Rat Tail Collagen I (Corning, 354236) prepared according to the manufacturer' s instruction. Human PDAC organoids were cultivated in growth factor reduced Matrigel$^{(R)}$ and overlaid in human complete medium [7]. Mouse organoids were established from normal pancreata, pancreatic tissues containing preinvasive lesions or full-blown carcinomas, and from metastatic tissues. Culture conditions for mouse pancreatic organoids are described in Hutch et al.[8]. Mouse PSC4 were kindly provided by Dr.

Tuveson (Cold Spring Harbor Laboratory), while NIH-3T3 were purchased from ATCC (CRL-1658).

Four of the five organoid models used were acquired as part of the Human Cancer Model Initiative (HCMI) https://ocg.cancer.gov/programs/HCMI and, those models are available for access from ATCC. The corresponding IDs are as follows:

| ID |
|---|
| HCM-CSHL-0080-C25 |
| HCM-CSHL-0077-C25 |
| HCM-CSHL-0081-C25 |
| HCM-CSHL-0092-C25 |

Both monolayer cell cultures and organoids were routinely tested for the presence of mycoplasma using MycoAlert Detection Kit from Lonza, in accordance with the manufacturer's instructions. The cultures were routinely cultured at 37°C with 5% $CO_2$. To grow cells in hypoxic conditions (5% or 3% $O_2$), cells were placed in an incubator that provides nitrogen in addition to $CO_2$ (Heracell 150i, ThermoFisher Scientific). Early-passage mouse tumor organoids were cultured in complete media supplemented with Nutlin-3a (10µM, Sigma-Aldrich) and propagated for at least 3 passages to generate cultures with bi-allelic alteration of p53 [9]. To induce epithelial-to-mesenchymal transition in organoids cultures, 5 ng/mL of TGF-$\beta$1 (Preprotech) was added to the culture media (depleted for A83-01 and mNoggin) for 5 days.

**Animal studies**

C57BL/6J and *NSG* (NOD.Cg-Prkdc$^{scid}$;Il2rg$^{tm1Wjl}$) mice were purchased from Charles River Laboratory (Milan). All animal experiments regarding transplanted mice were conducted in accordance with procedures approved by CIRSAL at University of Verona (approved project 655/2017-PR). For transplantation experiments, 6- to 8-week syngeneic C57BL/6J or immunodeficient NSG mice were used. For orthotopic transplantation, either cell lines (1x$10^5$ cells) or dissociated organoids (1x$10^6$ cells) were resuspended in 50 µL of a 2:3 dilution of Matrigel$^{(R)}$ (Corning) and cold PBS (Gibco) and injected into the tail region of the pancreas using insulin syringes (BD micro-fine 30 Gauge). The injection was considered successful by the development of bubble without signs of leakage. Monitoring of tumor growth was performed as previously described [10]. Briefly, following weekly manual palpation starting 7 days following transplantation, tumor-bearing mice were subjected to high-contrast ultrasound screening using the Vevo 2100 System with a MS250, 13–24 MHz scanhead (Visual Sonics). Mice were sacrificed at the indicated time points. Pancreas, spleen, lungs, and liver were collected for downstream analysis. For tail vein injection**,** cell lines (1x$10^5$ cells) were resuspended in 100 µL of PBS and injected in the lateral caudal vein using insulin syringes (BD micro-fine 30 Gauge). Mice were sacrificed at the indicated time points.

**Lentiviral infection of cell lines and organoids**

To knock out *Sema3a* in mouse PDAC organoids and cell lines, we used an inducible Crispr/Cas9 lentiviral system. First, both organoids and cell lines were transduced with an Edit-R Inducible Lentiviral hEF1a-Blast-Cas9 Nuclease Particles (Dharmacon, FE5VCAS11227) followed by selection with 5 µg/mL of blasticidin (Gibco). Three predesigned single-guide RNA (sgRNA) targeting *Sema3a,* and an individual sgRNA non-targeting control (NTC) were acquired from Dharmacon. For the transduction of monolayer cell cultures, 70-80% confluent cultures were incubated in complete transduction media (DMEM, 10% FBS, 1% P/S), lentiviral particles (MOI 10) and 1X polybrene (Santa Cruz Biotechnology); two days after the infection, 2 µg/mL of puromycin (Gibco) was added for selection. For organoids infection, cells were released from Matrigel® using 2mg/mL Dispase I for 20 minutes at 37°C. To obtain single cells, organoids were then enzymatically digested with TripLE (Gibco) supplemented with 2mg/mL Dispase I and 0.1 mg/mL DNAse I (Sigma-Aldrich) for 20 minutes at 37°C. $1x10^5$ cells were then incubated with transduction media (DMEM, 5% FBS, 1% P/S) supplemented with 1 µg/mL polybrene and lentiviral particles (MOI 10) and spinoculated for 1h at RT; cells were then incubated at 37°C for 16 hours and subsequently collected and embedded in 50 µL of Matrigel®. Two days after infection, cultures were treated with 2 µg/mL puromycin (Gibco) for antibiotic selection. Cas9 expression was induced by treating the cultures with 2.5 ng/mL of doxycycline for 3 days. Induction of Cas9 expression was evaluated by western blot using a mouse monoclonal antibody from Cell Signaling (clone 7A9-3A3, #14697). Successful editing and knock out were evaluated by Surveyor® Mutation Detection Kit (Integrated DNA Technologies) and western blotting, respectively. To overexpress *Sema3a* in mouse PDAC organoids and cell lines, we used a lentiviral vector carrying an open-reading frame for SEMA3A (tagged with MYC-DKK Origene, MR210565L3). LentiORF control particles of pLenti-C-Myc-DDK-P2A-Puro (Origene, PS100092V) were used as NTC, Lentiviral particles were packed in HEK293T cells transfected with plasmid containing the open-reading frame and the packaging plasmid VSV-G with X-tremeGENE9 (Roche, 063665110101). Lentiviral particles were collected two days after the infection by harvesting the viral surnatant, filtered and centrifuged to remove debris. Lentiviral particles were concentrated using Lenti-X concentrator (TaKaRa Bio) and viral titer was determined using Lenti-X qRT-PCR Titration Kit (TaKaRa Bio). Monolayer cell cultures transduction was performed by adding the viral supernatants supplemented with 1 µg/mL of Polybrene to cells having 50-60% of confluency. Antibiotic selection was started after 48 hours using 2 µg/mL of puromycin. Successful overexpression was assessed by Western blot. To knock down *Smad4* in mouse organoids, two shRNAs targeting *Smad4* were used. The vector TRC2 pLKO.5-puro (Sigma-Aldrich) was used as NTC. Organoids' infection was performed as described; two days after infection, cultures were treated with 2 µg/mL puromycin for antibiotic selection. Successful overexpression was assessed by western blot. To overexpress *Trp63* in mouse cultures, we used a lentiviral vector carrying an ORF for *Trp63* (mGFP-tagged; Origene, MR227530L4). Lentiviral particles production and monolayer cell cultures infection were performed as previously described. Successful overexpression was assessed by western blot.

**sgRNA targeting *Sema3a***

| sgRNA code | Sequence |
|---|---|
| 3A1 | GGAACTGCTGCGGACTTCAT |
| 3A2 | GTAAGGCACCCACTGATAGT |
| 3A3 | GATCAAGACCGGATATATGT |

**shRNA targeting *Smad4***

| shRNA code | Sequence |
|---|---|
| sh1 | TCAGGTAGGAGAGAGACGTTTAA |
| sh2 | GCGATTGTGCATTCTCAGGAT |

#### Histology and Immunohistochemistry

Both mouse and human tissues were fixed in 10% neutral buffered formalin and embedded in paraffin. Sections were subjected to Hematoxylin and Eosin as well as immunohistochemical staining. The following primary antibodies were used for immunohistochemical staining of human tissues: S100A2 [EPR5392] (109494, Abcam), CK5 [EP1601Y] (ab52635, Abcam), PDX1 [EPR3358(2)] (ab134150, Abcam), ΔNP63 (ACI3066C, Biocare), GATA-6 (AF1700, Bio-techne). The following primary antibody were used for immunohistochemical staining of mouse tissues: CD3e (SP7) (MA1-90582, Thermo Fisher), CD8a (4SM15) (14-0808-82, Invitrogen), F4/80 (ab6640, Abcam). Slides were scanned at 40x magnification and digitalized using the Aperio Scan-Scope XT Slide Scanner (Aperio Technologies). Quantification of CD3e, CD8a and F4/80 staining was performed in at least five random nonoverlapping fields of visualization (magnification, 20x) in each sample using ImageJ. To quantify CD3e and CD8a positive T cells, DAB positive cells were counted using the Aperio Image Scope software. To measure the percentage of positive F4/80 cells, captured images were first color deconvoluted and DAB+ particles counted automatically using ImageJ. Immunofluorescent staining of mouse tumor tissue was performed using Anti-mannose receptor antibody (CD206) (ab64693, Abcam) 1:100 and Pan-Keratin (C11) (#4545, Cell

Signaling Technology) 1:200. Slides were incubated with specific secondary antibodies at a 1:500 dilution: Goat anti-Rabbit IgG Alexa Fluor Plus 555 (A32732, Invitrogen) and Donkey anti-Mouse IgG Alexa fluor 488 (A21202, Invitrogen). Slides were then incubated with 0.1% Sudan Black B (Carlo Erba) in 70% ethanol for 20 min to reduce non-specific staining. Nuclei were stained using DAPI (D9542, Sigma-Aldrich). Images were captured with Leica TCS SP5 AOBS confocal system. Quantification was performed with ImageJ by measuring relative fluorescence of $CD206^+$ cells in at least three nonoverlapping fields.

**In situ hybridization**

The *in-situ* hybridization (ISH) was performed on 4 µm section of human PDAC tissues (n = 11). Briefly, sections were deparaffinized by incubation with xylene for 10 minutes, 100% ethanol for 2 minutes and then let dry for 5 minutes at room temperature. Slides were incubated for 10 minutes with RNAscope® Hydrogen Peroxide (Advanced Cell Diagnostics), washed with distilled water and incubated for 20 minutes at 99°C with RNAscope® 1X Retrieval Reagents (Advanced Cell Diagnostics). Sections were rinsed in distilled water and dehydrated in 100% ethanol for 3 minutes and let dry at room temperature. Then, the slides were incubated at 40°C for 10 minutes with RNAscope® Protease Plus (Advanced Cell Diagnostics), washed with distilled water and incubated with the appropriate probe for 2 hours at 40°C, followed by washes with RNAscope® 1X Wash Buffer (Advanced Cell Diagnostic). The different RNAscope® 2.5 HD AMPs (Hs-SEMA3A-C1 and Hs-PLXNA1-C2, Advanced Cell Diagnostics) were added to the slides following manufacturer's instructions. Positive control probe Hs-UBC and 2-plex negative control probe (Advanced Cell Diagnostics) were used as positive and negative control, respectively. Then, slides were incubated for 10 minutes at room temperature with either RNAscope® 2.5 HD Detection Reagent- RED or GREEN (Advanced Cell Diagnostic). Slides were stained with hematoxylin, dried at 60°C for 15 minutes and mounted with VectaMount® mounting medium (Vector Laboratories). Slides were scanned at 40x magnification and digitalized using the Aperio Scan-Scope XT Slide Scanner.

***In vitro* macrophages generation**

To isolate murine bone-marrow (BM) derived monocytes, we collected tibiae and femurs from 6 to 8-weeks old C57BL/6 mice and flushed BM cells. Red blood cells were lysed using a hypotonic solution (8.3% NH4Cl, 1% KHCO3 and 0.5M EDTA) [11]. $1{,}5*10^6$ BM derived cells were grown in RPMI 1640 (Euroclone) supplemented with 40 ng/mL IL-6 (Miltenyi Biotec, 130-096-686), 40 ng/mL GM-CSF (Miltenyi Biotec, 130-095-735), 10% FBS (Gibco), 2mM L-glutamine (Euroclone), 10mM HEPES (Euroclone), 1mM sodium pyruvate (Euroclone), 150U/mL streptomycin, 200U/mL penicillin/streptomycin. Cultures were maintained at 37°C in 5% $CO_2$-humidified atmosphere for 4 days. Macrophage's differentiation was promoted by growing BM derived cells in RPMI 1640 supplemented with 100ng/mL CSF-1 (Miltenyi Biotec, 130-101-706), 10% FBS, 2mM L-glutamine, 10mM HEPES, 1mM sodium pyruvate, 1% Pen/Strep [12]. Cells were grown for 7 days at 37°C in 5% $CO_2$-humidified atmosphere. On day 4 of the culture, fresh cytokine-supplemented complete medium was added. To obtain M1-MΨ or M2-MΨ, macrophages were cultured in presence of IFN-γ (10 U/mL; Miltenyi Biotec, 130-105-773) and LPS (1μg/mL; L8274; Sigma-Aldrich) or IL-4 (10 ng/mL; Miltenyi Biotec, 130-097-758) and IL-13 (10 ng/mL; Miltenyi Biotec, 130-096-688) cytokines-cocktail, respectively. Cells were either left untreated or treated with Recombinant Mouse Semaphorin 3A Fc (5926-S3-025/CF, Bio-techne, 100 ng/mL). To analyze macrophages polarization by flow cytometry, the following antibodies were used: anti-F4/80 FITC (Miltenyi Biotec, 130-117-509), anti-CD86 PE (Biolegend, 10-5106), anti-CD206 PerCP Cy5 (Biolegend, 141716), anti-CD80 APC (Biolegend, 104714). Cell viability was assessed using the LIVE/DEADFixable Aqua Dead Cell Stain Kit (Invitrogen, L34966).

**_In vitro_ macrophages polarization**

Raw 264.7 cells were grown in DMEM supplemented with 10% FBS, 1% pen/strep, 2 mmol/L l-glutamine, 10 mmol/L HEPES. To induce *in vitro* differentiation, cells were stimulated with M-CSF (25 ng/ml) for 48hs, followed by a 72h incubation with IL4 (40 ng/ml) and IL13 (40 ng/ml) to promote M2 polarization or conditioned medium collected from FC1199 CTR or FC1199 3A-OE cell lines. Unstimulated RAW 264.7 (M0) were used as control.

**Transwell migration assay**

To assess the chemo-attractant capability of SEMA3A towards macrophages, we performed a transwell migration assay using the cell line RAW264.7 (kindly provided by Prof. Vincenzo Bronte, University of Verona). Cells were grown in DMEM supplemented with 10% FBS, 1% pen/strep, 2 mmol/L l-glutamine, 10 mmol/L HEPES and differentiated into M1 or M2 macrophages. To obtain M1 macrophages, cells were stimulated with M-CSF (25 ng/ml) for 48 h, followed by a 72-hour stimulation with IFN γ (0.3 μg/ml) and LPS (1μg/ml). For M2 macrophages polarization, M-CSF stimulation was followed by a 72 h IL4 (40 ng/ml) and IL13 (40 ng/ml) stimulation. Non-stimulated (M0) cells were used as control. To perform the *in vitro* migration assay, we used 8 μm pore transwell (Corning, 353097) coated with 10% Matrigel. Both M1 and M2 polarized RAW264.7, and unstimulated cells, were collected and resuspended in 2% FBS growth medium and seeded on top of the coated transwell [13]. Growth medium with or without 100 ng/ml SEMA3A-Fc was added on the bottom of the well. Growth medium supplemented with 20% FBS was used as positive control. After 24 h, non-migrated cells were removed from the upper side of the transwell with a cotton swab whereas transwells' membranes were fixed with 3.7% Paraformaldehyde (Sigma Aldrich, 8187081000), stained with crystal violet and eluted with a solution containing 50% ethanol and 0.1% acetic acid. Absorbance was measured at 595 nm using a microplate reader (BioTek, Synergy 2 Multi-mode Microplate Reader).

#### *In vivo* treatment experiment

For *in vivo* treatment with anti-CSF1R, 7-8 weeks old C57BL6/J mice of both sexes were randomized into four groups and treated for 3 days with 300 µg of anti-mouse CSF1R (CD115) [AFS98] (BioXCell, BE0213) or rat IgG2a [2A3] (BioXCell, BE0089) as isotypic control. On day four, mice were orthotopically transplanted with $1*10^5$ FC1199 NTC or FC1199 3A-OE cells. Animals were treated every 2 days and tumor growth was monitored by abdominal palpation and high-contrast ultrasound (Vevo 2100 System). Mice were sacrificed when tumors reached an average volume of approximately 100 $mm^3$. To evaluate the effect of anti-CSF1R treatment on immune cell populations, the peripheral blood from untreated and treated mice was subjected to cytofluorimetric analysis with the following antibodies: anti-CD45.2 APC (Invitrogen, 14-0454-82), anti-CD11b FITC (M1/70) (eBioscience, 11-0112-85), anti-F4/80 PECy7 (BM8) (eBioscience, 25-4801-82), anti-Ly6C V450 (Invitrogen, 48-5932-82), anti-Ly6G APC-H7 (Biolegend, 127624), anti-CD16/32 (Sony,1106510). Cell viability was assessed using the LIVE/ DEADFixable Aqua Dead Cell Stain Kit. Samples were acquired with FACS-Canto II (BD). Data were analyzed by FlowJo software (Tree Star, Inc). Survival study was performed using 7-8 weeks old female C57BL6/J mice orthotopically transplanted with $1*10^5$ FC1199 NTC or FC1199 3A-OE cells. On day 10, tumor growth was assessed by abdominal palpation and mice were randomized into 8 groups. Mice were treated 3 times/week with 300 µg of anti-mouse CSF1R (CD115) [AFS98], rat IgG2a [2A3] as isotypic control or 120 mg/kg Gemcitabine once a week.

#### Transient Knockdown in Cell lines

To transiently downregulate the expression of specific genes, cells were transfected with siRNA for *Kras*, *TP63*, or with scramble control siRNA, and harvested 48 hours later. Mouse cell lines were transfected with 5 nM of ON-TARGET Plus mouse Kras siRNA, SMART pool (horizon, L-043846-01-0005). Human cell lines were transfected with 25 pmol of siRNA targeting *TP63* (Thermofisher Scientific, 4392420, Assay ID: s531582). Lipofectamine™ 2000 (Thermofisher Scientific, #11668027) was used for siRNA transfection according to manufacturer's instruction for 6-well plates format.

#### Cell viability assay

The proliferation of mouse cells following genetic perturbation of *Sema3a* was measured using the CellTiter-Glo assay (Promega, G9683). $1*10^3$ cells were plated on white 96-well plate (Thermofisher Scientific), cultured in 100 µL of culture medium, and viability was measured at endpoint (72 hours) using a microplate reader (BioTek, Synergy 2 Multi-mode Microplate Reader).

**Immunoblotting**

Whole cell lysates were prepared using Cell Signaling Lysis Buffer (Cell Signaling) and separated on 4-12% Bis-Tris NuPAGE gels (Life Technologies), transferred onto a PVDF membrane (Thermo Scientific) and incubated with the following antibodies: p63 (D9L7L) (39692, Cell Signalling), Sema3a (ab23393, Abcam), Sema3c (MAB1728, bio-techne) Smad4 (B-8/sc-7966, Santa Cruz Biotechnology), phospho-FAK (Tyr-397) (31H5L17/700255, Invitrogen), FAK (3285, Cell Signalling), phospho-Akt (4060, Cell Signalling), Akt (9272, Cell Signalling), ERK1/2 (9102, Cell Signalling), phospho-ERK1/2 (4376, Cell Signalling), E-Cadherin (M3612, Dako), Vimentin (NCL-L-VIM-V9, Leica Biosystems). GADPH (5174, Cell Signalling) (sc-166545, Santa Cruz Biotechnology) and β-Actin (4967, Cell Signalling) were used as loading control. The immunoblots were visualized with ECL plus (Amersham/GE Healthcare Europe GmgH).

**Anoikis and Scratch assays**

The anoikis assay was performed on both single cells dissociated from mouse metastatic organoids deficient for *Sema3a* and mouse monolayer cell cultures (FC1199) overexpressing *Sema3a.* Cells were plated in Poly(2-hydroxyethyl methacrylate) (sc-253284) coated 96-well plates black/clear flat tc (Corning) ($5*0^3$ cells/well for mouse organoids; $1*10^3$ cells/well for FC1199). Cells were either left untreated or treated with Y-27632 dihydrochloride (Y0503, Sigma-Aldrich, 10,5 µM), Recombinant Mouse Semaphorin 3A Fc (100 ng/mL), or the selective FAK inhibitor Defactinib (Selleckem, 1µM). CellEvent™ Caspase-3/7 Green Detection Reagent (Invitrogen) was added to the cultures at a concentration of 4 µM. After 4 days, Hoechst (1µg/ml, Thermo Fisher) was added to the cultures and incubated for 30 minutes. Images were acquired using the EVOS Cell Imaging System (Thermo Fisher Scientific) and green signal quantification was performed in at least five random nonoverlapping fields using ImageJ. For the scratch assay, FC1199 3A3-KO, FC1199 3A-OE and FC1245 3A-OE monolayer cell cultures were initially plated in DMEM, 10% FBS, 1% P/S in 6-well plates. Once cells reached 80% confluency, the culture medium was replaced with a low-serum medium (2% FBS) and cells cultivated in this condition for additional 24 hours. A 200 µl pipette tip was then used to scrape the cells followed by two consecutive washes with PBS to remove cell debris. Finally, fresh low-serum culture medium was added to the cultures. Recombinant Mouse Semaphorin 3A Fc was used at a final concentration of 100 ng/mL. Brightfield images were taken at 0h, 8h and 24h at a 2.5X magnification.

**qRT-PCR analysis**

RNA was extracted from cell lines and organoids using the Trizol® Reagent (Life Technologies) method. 1 µg of DNase-treated RNA was reverse transcribed using

TaqMan® Reverse Transcription reagents (Applied Biosystems) in a volume of 20 µL according to the manufacturer's instructions. Samples were diluted to a final concentration of 10 ng/µL. For TaqMan-based qRT-PCR. the following probes were used (TaqMan® Gene Expression Assay): *Sema3a* (Mm00436469_m1), *Ctgf* (Mm01192933_g1 *),* *Zeb1* (Mm00495564_m1), *Snai* (Mm00441533_g1), *Trp63* (Mm00495793_m1), *Smad4* (Mm03023996_m1), *Itga2* (Mm00434371_m1), *Itga1* (Mm01306375_m1), *Itgb1* (Mm01253230_m1), *Vegf (*Mm01281449_m1), *Kras* (Mm00517492_m1), *Sema3c* (Mm00443121_m1), *Gata6* (Mm00802636_m1), *Nqo1* (Mm01253561_m1), *Rpa3* (Mm01165241_g1), TP63 (Hs00978340_m1), SEMA3A (Hs00173810_m1), Hprt1/HPRT1 (Mm03024075_m1, Hs02800695_m1) was used as reference gene was used as reference gene. For SYBR green-based qRT-PCR the following primers were used: *Inos* (F: GTTCTCAGCCCAACAATACAAGA, R: GTGGACGGGTCGATGTCAC), *Arg1* (F: CTCCAAGCCAAAGTCCTTAGAG, R: AGGAGCTGTCATTAGGGACATC), *Fizz1* (F: TCCCAGTGAATACTGATGAGA, R: CCACTCTGGATCTCCCAAGA), *Hprt1* (F: CTGGTGAAAAGGACCTCTCGAAG, R: CCAGTTTCACTAATGACACAAACG). Relative gene expression quantification was performed using the ΔΔCt method with the Sequence Detection Systems Software, Version 1.9.1 (Applied Biosystems).

**Chip qPCR**

Mouse cell lines overexpressing *Trp63* (1X$10^7$ cells) were fixed for 10 minutes with 37% formaldehyde, while slowly shaking. Formaldehyde was quenched with 125 mM glycine; medium was removed, and cells were washed with cold PBS. Cells were collected and centrifuged at 1500 rpm for 10 minutes at 4°C. Cell pellet was resuspended in cell lysis buffer (50 mM Tris, pH 8.0, 140 mM NaCl, 1mM EDTA, 10% glycerol, 0,5% NP40, 0,25% Triton X-100) and incubated for 10 minutes on ice. Cell nuclei were pelleted at 1000 rpm (10 minutes, 4°C) and then resuspended in Nuclei lysis buffer (10 mM Tris pH 8.0, 1mM EDTA, 0,5 mM EGTA, 0,2% SDS), supplemented with protease inhibitors (cOmplete(TM), Mini Protease Inhibitor, Sigma-Aldrich), and incubated for 10 minutes on ice before proceeding to sonication. Nuclei were sonicated by six 15-seconds pulses followed by 45-seconds rest periods (50% maximum potency). Chromatin was then transferred to a 1,5 mL microcentrifuge tube and centrifuged at 8000 rpm for 15 minutes at 4°C. Supernatant was collected for chromatin immunoprecipitation and DNA fragment size evaluation. DNA was extracted using DNeasy Blood & Tissue Kit (Qiagen) and DNA concentration evaluated with Nanodrop 2000 (Life Technologies) and 100 mg of chromatin was used for each reaction. To perform chromatin immunoprecipitation ChIP grade protein A/G plus agarose (50% slurry) was used (Life technologies); chromatin was diluted to a final volume of 300 mL in dilution buffer (10 mM Tris pH 8.0, 0,5 mM EGTA, 1% Triton X-100, 140 mM NaCl) supplemented with protease inhibitors. To preclear chromatin, samples were incubated with 10 mL of protein A/G for 1h at 4°C;

samples were then centrifuged at 3000 rpm for 5 minutes at 4°C and the surnatant transferred to a fresh microcentrifuge tube, incubated for 1h at 4°C with p63 (D9L7L) antibody (Cell Signaling Technologies) or normal rabbit IgG (2729S, Cell Signaling Technologies) and then incubated over night at 4°C with 50 mL of protein A/G.
To elute immunoprecipitated chromatin, samples were centrifuged for 1 min at 3000 rpm, surnatant was removed and samples were subjected to 4 10 minutes incubation with 1 mL of high-salt wash buffer (2mM EDTA, 20 mM HEPES pH 8.0, 500 mM NaCl, 0,1% SDS, 1% Triton X-100) was added to each sample; beads were then incubated with 300 mL of elution buffer (50 mM Tris pH 8.0, 10 mM EDTA, 1% SDS) supplemented with Proteinase K (20 µg/µL), incubated for 2h at 55°C and then overnight at 65°C to reverse crosslink. Immunoprecipitated chromatin was centrifuged for 5 minutes at room temperature, subjected to purification, and resuspended in Nuclease-free Water (Invitrogen) for qPCR analysis.
For qPCR analysis, 7 different loci of the *Sema3a* promoter were targeted (see table below); *Krt5* was used as a positive control for the immunoprecipitation whereas normal rabbit IgG were used as negative control; *Hprt1* was used as housekeeping gene.

| **Locus name** | **Primers sequence** |
|---|---|
| *Sema3a* 1 | F: ACAAGCCAACTGTGAGAACA<br>R: GATATGCAATGGTGAGGTGGG |
| *Sema3a* 2 | F: TGTCCAATCACAGATCCCCT<br>R: ACCAACCACACTACCAGCAT |
| *Sema3a* 3 | F: TCTTGAGGACAGACTGAACCC<br>R: TCATCTCTGTTGTGTCCTGCT |
| *Sema3a* 4 | F: AGCTCTCTTCGACATAAGGCA<br>R: TGCTCCTCATTATACGGGGT |
| *Sema3a* 5 | F: AGCTCTCTTCGACATAAGGCA<br>R: CTGACACTCTGCAATTGGGC |
| *Sema3a* 6 | F: TTAACGCTACCCCACTTCCA<br>R: TGCCGTTGTCACACTTGTTT |
| *Sema3a* 7 | F: CGTTTGTGCCTAACCCAGAG<br>R: CACAGGCTTCTTTTGCTGGT |
| *Krt5* 1 | F: TACCCAAGAAAAGAGGCCGT<br>R: CAGCTTTGGGGTTTCTGTCC |
| *Krt5* 2 | F: CCAGGACACCAGCTCTGTAA<br>R: TAGAAGTGGGTTGGGCAGAG |
| *Krt5* 3 | F: ATCCTACAGTTCTGGCGGAG<br>R: CGATTTGCAGGTTCAGAGGG |

| | |
|---|---|
| *Hprt1* | F: CTGGTGAAAAGGACCTCTCGAAG<br>R: CCAGTTTCACTAATGACACAAACG |

**ELISA**

Cell culture supernatants were collected after 3 days of conditioning; particulates were removed by centrifuging samples for 15 minutes at 1000g, 4°C. Plasma from orthotopic mouse models was collected using EDTA 0,5 M as an anticoagulant and centrifuged for 15 minutes at 1000g, 4°C within 30 minutes of collection. ELISA assay was performed using Mouse Semaphorin-3A(SEMA3A) ELISA kit (CUSABIO, CSB-EL020980MO-96) following manufacturer's instructions. Results were calculated using the Curve Expert 1.4 software.

**RNA sequencing**

RNA was harvested from monolayer cell cultures using TRIzol (LifeTechnologies), followed by column-based purification with the PureLink RNA Mini Kit (Ambion). To collect RNA from flash-frozen tumor tissues, at least 30 consecutive cryosections (16 µm) for each sample were cut and collected in TRIzol. RNA isolation was performed using the PureLink RNA Mini Kit. The quality of purified RNA was evaluated using a Bioanalyzer 2100 (Agilent) with an RNA 6000 Nano Kit. RNAs with RNA Integrity Number (RIN) values greater than 8 were used for the generation of sequencing libraries using the TruSeq sample Prep Kit V2 (Illumina) according to the manufacturer's instructions. RNA-Seq libraries prepared from in vitro cultures (n = 12) and from in mouse PDAC tissues (n = 5) were multiplexed and sequenced using a NextSeq 500 platform with single-end reads of 75 bases and a final coverage of 3 million reads per sample. First, we performed quality control and trimming steps, then we aligned reads to the mm10 genome build using Salmon v1.4.0 [14]. Then, we quantified transcripts using the R package tximport v4.0 [15]. Finally, we normalized count data with the R/ Biocondutor package DESeq2 v1.30.0 [16], which was also used to identify differentially expressed genes. For gene set enrichment analysis (GSEA), we used the fgsea R package v1.16.0 [17] based on the list of differentially expressed genes sorted by log2 of fold change. We retrieved pathways from MSigDB database and in particular Gene Ontology, KEGG, Biocarta, Reactome and Hallmark gene sets. fgsea function was used with default parameters. Results were considered significant for FDR < 0.05. To classify PDAC cultures and tissues according to the main transcriptomic subtypes [1, 3], we calculated the score for each of the gene sets (i.e., basal-like, classical, pancreatic progenitor, squamous, gene program 2 and gene program 3) using the GSVA R package v1.38.2 [18]. The same package was used to evaluate the enrichment of custom defined immune cell signatures. In both cases, Gsva function was used with ssgsea and gene set scores were compared among different experimental conditions with Wilcoxon rank-sum test.

**Analysis of Single-cell RNA-Sequencing data**

scRNA-Seq data from normal pancreatic tissues were downloaded from the Seurat V3 repository together with their annotation metadata. The four datasets Muraro et al. [19] (ncells=2285), Segerstolpe et al.[20] (cells = 2394), Grün et al.[21] (ncells = 1004) and Lawlor et al. [22] (ncells = 638) were integrated and queried with the *R* package Seurat V4.0.1 [23]. ScRNA-Seq from human PDAC tissues PDAC tissues were downloaded from NGDC (GSA: CRA001160), GEO (Accession #GSE154778 and #GSE155698) and EGA (accession EGAS00001002543) together with annotation metadata, when provided. The dataset Peng et al.[5] (primary PDAC = 24, ncells = 41964), Lin et al.[24] (primary PDAC = 10, ncells = 7752), Chan-Seng-Yue et al.[4] (primary PDAC = 13, ncells = 33970) and Steele et al.[25] (primary PDAC = 16, ncells = 42844) were first preprocessed individually using Seurat V4.0.1[23] for quality control and filtering (percent_mt_max = 20, nFeature_min = 500, nCount_min = 500, nCount_max = 50000), then integration was performed through *harmony* [26] using default parameters and dataset metadata as grouping variable. The integrated dataset was annotated through *singleR* package using as reference the preloaded dataset *HPCA* from the *celldex* package [27].

**Analysis of DNA Methylation datasets**

The data relative to the correlation between methylation level and SEMA3A expression in pancreatic progenitor and squamous PDAC of the ICGC cohort were retrieved from the supplementary tables of the original publication [3]. The inspected probes were the following (reference genome GRCh37/hg19):

| | |
|---|---|
| cg11493571 | intron 9 |
| cg10120616 | intron 1 |
| cg06144675 | promoter |
| cg05081033 | promoter |

**Statistical Analysis and datamining**

We analyzed three available transcriptomic datasets to explore the correlation between SEMA3A expression and PDAC molecular subtypes. Those cohorts are: the PACA-AU cohort [3] of the ICGC consortium; the TCGA-PAAD cohort [2] of the TCGA consortium and the PanCuRx cohort [4] (EGA archive accession EGAS00001002543). The ICGC dataset contains normalized expression values (TMM normalized using edgeR Bioconductor package, converted to CPM and log2 transformed) of 96 pancreatic cancer patients. We downloaded associated clinical data from https://dcc.icgc.org/releases/current/Projects/PACA-AU. The TCGA dataset was downloaded from http://

firebrowse.org/?cohort=PAAD, and the sample number filtered down to 148, according to a more precise histological revision of included specimens. The PanCuRx dataset was preprocessed with STAR v2.7.6a [28] and RSEM v1.3.1 [29], and eventually a vst expression matrix was produced. Within each cohort, Z-scores standardization of the expression data was performed before subsequent analysis. Expression of SEMA3A was stratified according to tumor stage, subtype classification, and (when available) survival status. The correlation of SEMA3A expression with other genes was evaluated using Spearman's correlation test (significant p-value $< 0.001$). To classify tumors from the PanCuRx and the ICGC datasets according to the expression of SEMA3A, the distribution of their transcripts levels in the respective cohorts was assessed. We observed a normal distribution for ICGC and TCGA, whereas PanCuRx was bimodal. Thus, we stratified the first two cohorts according to median value of SEMA3A expression. In the PanCuRx dataset instead, we set a threshold at the 3$^{rd}$ quartile which corresponded to 5 (vst).

### REFERENCES TO SUPPLEMENTAL MATERIAL


1 Moffitt RA, Marayati R, Flate EL, Volmar KE, Loeza SG, Hoadley KA*, et al.* Virtual microdissection identifies distinct tumor- and stroma-specific subtypes of pancreatic ductal adenocarcinoma. Nat Genet 2015;**47**:1168-78.

2 Cancer Genome Atlas Research Network. Electronic address aadhe, Cancer Genome Atlas Research N. Integrated Genomic Characterization of Pancreatic Ductal Adenocarcinoma. Cancer Cell 2017;**32**:185-203 e13.

3 Bailey P, Chang DK, Nones K, Johns AL, Patch AM, Gingras MC*, et al.* Genomic analyses identify molecular subtypes of pancreatic cancer. Nature 2016;**531**:47-52.

4 Chan-Seng-Yue M, Kim JC, Wilson GW, Ng K, Figueroa EF, O'Kane GM*, et al.* Transcription phenotypes of pancreatic cancer are driven by genomic events during tumor evolution. Nat Genet 2020;**52**:231-40.

5 Peng J, Sun BF, Chen CY, Zhou JY, Chen YS, Chen H*, et al.* Single-cell RNA-seq highlights intra-tumoral heterogeneity and malignant progression in pancreatic ductal adenocarcinoma. Cell Res 2019;**29**:725-38.

6 Hingorani SR, Wang L, Multani AS, Combs C, Deramaudt TB, Hruban RH*, et al.* Trp53R172H and KrasG12D cooperate to promote chromosomal instability and widely metastatic pancreatic ductal adenocarcinoma in mice. Cancer Cell 2005;**7**:469-83.

7 Boj SF, Hwang CI, Baker LA, Chio, II, Engle DD, Corbo V*, et al.* Organoid models of human and mouse ductal pancreatic cancer. Cell 2015;**160**:324-38.

8 Huch M, Bonfanti P, Boj SF, Sato T, Loomans CJ, van de Wetering M*, et al.* Unlimited in vitro expansion of adult bi-potent pancreas progenitors through the Lgr5/R-spondin axis. EMBO J 2013;**32**:2708-21.

9 Oni TE, Biffi G, Baker LA, Hao Y, Tonelli C, Somerville TDD*, et al.* SOAT1 promotes mevalonate pathway dependency in pancreatic cancer. J Exp Med 2020;**217**.

10 Filippini D, Agosto S, Delfino P, Simbolo M, Piro G, Rusev B*, et al.* Immunoevolution of mouse pancreatic organoid isografts from preinvasive to metastatic disease. Sci Rep 2019;**9**:12286.

11 Facciabene A, De Sanctis F, Pierini S, Reis ES, Balint K, Facciponte J*, et al.* Local endothelial complement activation reverses endothelial quiescence, enabling t-cell homing, and tumor control during t-cell immunotherapy. Oncoimmunology 2017;**6**:e1326442.

12 Solito S, Pinton L, De Sanctis F, Ugel S, Bronte V, Mandruzzato S*, et al.* Methods to Measure MDSC Immune Suppressive Activity In Vitro and In Vivo. Curr Protoc Immunol 2019;**124**:e61.

13 Marigo I, Trovato R, Hofer F, Ingangi V, Desantis G, Leone K*, et al.* Disabled Homolog 2 Controls Prometastatic Activity of Tumor-Associated Macrophages. Cancer Discov 2020;**10**:1758-73.

14 Patro R, Duggal G, Love MI, Irizarry RA, Kingsford C. Salmon provides fast and bias-aware quantification of transcript expression. Nat Methods 2017;**14**:417-9.

15 Soneson C, Love MI, Robinson MD. Differential analyses for RNA-seq: transcript-level estimates improve gene-level inferences. F1000Res 2015;**4**:1521.

16 Love MI, Huber W, Anders S. Moderated estimation of fold change and dispersion for RNA-seq data with DESeq2. Genome Biol 2014;**15**:550.
17 Subramanian A, Tamayo P, Mootha VK, Mukherjee S, Ebert BL, Gillette MA*, et al.* Gene set enrichment analysis: a knowledge-based approach for interpreting genome-wide expression profiles. Proc Natl Acad Sci U S A 2005;**102**:15545-50.
18 Hanzelmann S, Castelo R, Guinney J. GSVA: gene set variation analysis for microarray and RNA-seq data. BMC Bioinformatics 2013;**14**:7.
19 Muraro MJ, Dharmadhikari G, Grun D, Groen N, Dielen T, Jansen E*, et al.* A Single-Cell Transcriptome Atlas of the Human Pancreas. Cell Syst 2016;**3**:385-94 e3.
20 Segerstolpe A, Palasantza A, Eliasson P, Andersson EM, Andreasson AC, Sun X*, et al.* Single-Cell Transcriptome Profiling of Human Pancreatic Islets in Health and Type 2 Diabetes. Cell Metab 2016;**24**:593-607.
21 Grun D, Muraro MJ, Boisset JC, Wiebrands K, Lyubimova A, Dharmadhikari G*, et al.* De Novo Prediction of Stem Cell Identity using Single-Cell Transcriptome Data. Cell Stem Cell 2016;**19**:266-77.
22 Lawlor N, George J, Bolisetty M, Kursawe R, Sun L, Sivakamasundari V*, et al.* Single-cell transcriptomes identify human islet cell signatures and reveal cell-type-specific expression changes in type 2 diabetes. Genome Res 2017;**27**:208-22.
23 Hao Y, Hao S, Andersen-Nissen E, Mauck WM, 3rd, Zheng S, Butler A*, et al.* Integrated analysis of multimodal single-cell data. Cell 2021;**184**:3573-87 e29.
24 Lin W, Noel P, Borazanci EH, Lee J, Amini A, Han IW*, et al.* Single-cell transcriptome analysis of tumor and stromal compartments of pancreatic ductal adenocarcinoma primary tumors and metastatic lesions. Genome Med 2020;**12**:80.
25 Steele NG, Carpenter ES, Kemp SB, Sirihorachai V, The S, Delrosario L*, et al.* Multimodal Mapping of the Tumor and Peripheral Blood Immune Landscape in Human Pancreatic Cancer. Nat Cancer 2020;**1**:1097-112.
26 Korsunsky I, Millard N, Fan J, Slowikowski K, Zhang F, Wei K*, et al.* Fast, sensitive and accurate integration of single-cell data with Harmony. Nat Methods 2019;**16**:1289-96.
27 Aran D, Looney AP, Liu L, Wu E, Fong V, Hsu A*, et al.* Reference-based analysis of lung single-cell sequencing reveals a transitional profibrotic macrophage. Nat Immunol 2019;**20**:163-72.
28 Dobin A, Davis CA, Schlesinger F, Drenkow J, Zaleski C, Jha S*, et al.* STAR: ultrafast universal RNA-seq aligner. Bioinformatics 2013;**29**:15-21.
29 Li B, Dewey CN. RSEM: accurate transcript quantification from RNA-Seq data with or without a reference genome. BMC Bioinformatics 2011;**12**:323.
